# Exploiting correlations across trials and behavioral sessions to improve neural decoding

**DOI:** 10.1101/2024.09.14.613047

**Authors:** Yizi Zhang, Hanrui Lyu, Cole Hurwitz, Shuqi Wang, Charles Findling, Yanchen Wang, Felix Hubert, International Brain Laboratory, Alexandre Pouget, Erdem Varol, Liam Paninski

## Abstract

Traditional neural decoders model the relationship between neural activity and behavior within individual trials of a single experimental session, neglecting correlations across trials and sessions. However, animals exhibit similar neural activities when performing the same behavioral task, and their behaviors are influenced by past experiences from previous trials. To exploit these informative correlations in large datasets, we introduce two complementary models: a multi-session reduced-rank regression model that shares similar behaviorally-relevant statistical structure in neural activity across sessions to improve decoding, and a multi-session state-space model that shares similar behavioral statistical structure across trials and sessions. Applied across 433 sessions spanning 270 brain regions in the International Brain Laboratory public mouse Neuropixels dataset, our decoders demonstrate improved decoding accuracy for four distinct behaviors compared to traditional approaches. These results generalize across additional datasets, species, and behavioral tasks. Unlike existing deep learning approaches, our models are interpretable and efficient, uncovering low-dimensional representations that predict animal decisions, quantifying single-neuron contributions to decoding behaviors, and identifying different activation timescales of neural activity across the brain. Code: https://github.com/yzhang511/neural_decoding.

## 1 Introduction

Neural decoding is a critical tool for understanding the relationship between behavior and brain activity. Traditional neural decoders operate within a single-trial, single-session context (Paninski et al., 2007; Glaser et al., 2020), modeling the relationship between neural activity and behavior within individual trials of each experimental session. However, these decoders overlook informative correlations across trials and sessions in both the neural and behavioral data, missing opportunities to leverage information from large datasets collected across numerous experiments.

Similar neural activities emerge across experimental sessions when animals engage in the same behavioral task (Gallego et al., 2018; Melbaum et al., 2022; Pellegrino et al., 2023; Safaie et al., 2023). Incorporating such inter-session neural similarities offers an opportunity to improve single-session decoding accuracy (Herrero-Vidal et al., 2021; Schneider et al., 2023; Chen et al., 2021). However, directly sharing this information across sessions is challenging, since typically different populations of neurons are recorded in each session. An alternative approach is to focus on the important neural population variations relevant to the behavior, utilizing their correlation structures across sessions. Previous unsupervised studies have adopted this strategy to improve neural dynamics estimation by sharing activities across sessions (Turaga et al., 2013; Pandarinath et al., 2018; Ye et al., 2023). However, the learned neural latents may not be behaviorally relevant, and have to be fine-tuned for supervised decoding tasks. While supervised and self-supervised pre-training can learn shared neural representations by training models on multiple sessions before fine-tuning them to decode specific behaviors, existing methods (Azabou et al., 2023; Zhang et al., 2025) require substantial computing resources and result in complex black-box models that lack interpretability. For a more lightweight and interpretable solution, a simple yet effective model is needed for sharing behaviorally relevant neural variations across many sessions.

Similarly, animal behavior is shaped not only by the current task, but also by the animal’s experiences from previous trials. For example, Ashwood et al. (2022) found that mouse decision-making shows internal states persisting across tens to hundreds of trials, effectively modeled by hidden Markov models (HMMs). These latent states are reproducible across animals and experiment sessions. Many neuroscience experiments exhibit trial-to-trial behavioral correlations arising from such reproducible latent states. Explicitly accounting for these behavioral correlations across sequential trials, in addition to modeling intersession neural similarities, can potentially improve neural decoding performance. In addition, inferring latent states from trial-to-trial correlations allows us to gain insight into the internal states driving the animal’s behavior, by observing how animals transition between states during a task. Investigating whether these latent states are shared across individuals can reveal whether such internal states reflect a common behavioral strategy or are specific to each animal.

In this work we develop two complementary methods to leverage these neural and behavioral correlations for improved neural decoding. For neural data, we employ a multi-session reduced-rank regression model that shares similar temporal patterns in the neural activity across sessions while retaining session-specific differences to accommodate individual variations. For behavioral data, we use multi-session state-space models to learn latent behavioral states from trial-to-trial correlations in animal behaviors across multiple sessions. These learned neural and behavioral representations are then used to improve single-trial, single-session decoders. Unlike existing deep learning methods that share data across sessions through complex black-box models, our models are simple, highly interpretable, and easy to fit. We evaluate our neural and behavioral data-sharing models using mouse Neuropixels recordings from the International Brain Lab (IBL et al., 2022, 2023), which include 433 sessions and 270 brain regions. We additionally apply our models to the Allen Institute Neuropixels visual coding dataset (de Vries et al., 2023) and a primate random target reaching task with Utah array recordings (Pei et al., 2021). The results demonstrate improved decoding accuracy across datasets, species, and behavioral tasks. Our approach is computationally efficient and enables us to create a brain-wide map of behaviorally-relevant timescales and identify key neurons associated with each behavioral task.

## 2 Formulation of the neural data-sharing model

The goal of our analysis is to decode the behavior *y* ∈ ℝ^*P*^ from spike-sorted and temporally-binned spike count data. Given neural activity of size *N* × *T* from each trial indexed by *k*, where *N* is the number of neurons and *T* is the number of timesteps, we aim to decode the behavior per trial. In most of the analyses presented here, we split the recording into equal-length trials of 2 seconds. We further divide each trial into 20-millisecond time bins, yielding 100 timesteps per trial. For each trial within a session, we construct the input *X* ∈ ℝ^*N*×*T*^ by aggregating the spike counts of *N* neurons over *T* = 100 timesteps, and use it to compute a decoder estimate *d* ∈ ℝ^*P*^ of the true behavior *y* ∈ ℝ^*P*^. When *P* = 1, the value of *y* remains constant throughout a trial (*per-trial decoded behavior*). When *P* = *T*, the value of *y* varies over time within a trial (*per-timestep decoded behavior*). To simplify notation, we initially present the model assuming *y* is a scalar (i.e., *P* = 1).

Traditional single-session decoders often rely on *full-rank* models, which predict the output using a full-rank (unconstrained) weight matrix *W* ∈ ℝ^*N*×*T*^ :

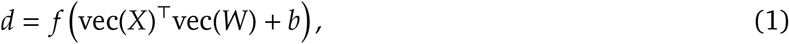

where vec(·) denotes the vectorization (flattening) operator and *b* ∈ ℝ is a bias term. However, when the number of neurons *N* and time steps *T* is large, this approach becomes prone to overfitting due to the large number of parameters.

To address this, we introduce a low-rank constraint on the weight matrix by factorizing it into two smaller matrices:

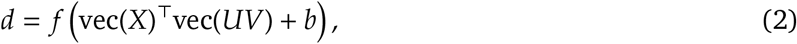

where *U* ∈ ℝ^*N*×*R*^ and *V* ∈ ℝ^*R*×*T*^ are low-rank bases that represent the neural and temporal components of the original weight matrix *W*. This factorization, referred to as a **reduced-rank regression model (RRR)**, reduces the number of parameters from *N* × *T* in the full-rank model (Equation 1) to *R*×(*N* + *T*), where *R* < min(*N, T*). The function *f* can be linear or nonlinear, depending on the application. Note that the reduced-rank regression model proposed in Equation 2 differs from the standard reduced-rank regression that has previously been applied to predict neural responses (Posani et al., 2025); we elaborate on these differences in the STAR Methods section.

The solutions of *U, V* and *b* can be obtained through either automatic differentiation or closed-form expressions. When *f* is an identity function (linear model), closed-form solutions are attainable. The closed-form solution of *U* reveals that it can be interpreted as a subspace that maximizes the correlation between neural activity *X* and behavior *y* while capturing the major variations in *y*. Thus, this reduced-rank regression model can be viewed as a latent variable model, where the rank *R* determines the number of latent variables required to capture the behaviorally relevant variations in neural activity. See STAR Methods, section “Closed-form solution for reduced-rank regression model” for details.

Instead of manually aligning neurons from different populations based on their firing or physical properties (Haxby et al., 2020; Busch et al., 2021), we aim to automatically learn a common neural representational space crucial for decoding from multiple neural populations. To this end, we introduce a **multi-session reduced-rank regression model** (Figure 1C) to learn such common neural representations and improve neural decoding. Since neural populations within a given region may exhibit similar activation patterns (Gallego et al., 2018) (Figure 1B), we can share the low-rank temporal basis set *V* across sessions and retain session-specific differences via the neural basis set *U*^*i*^:

**Figure 1:**
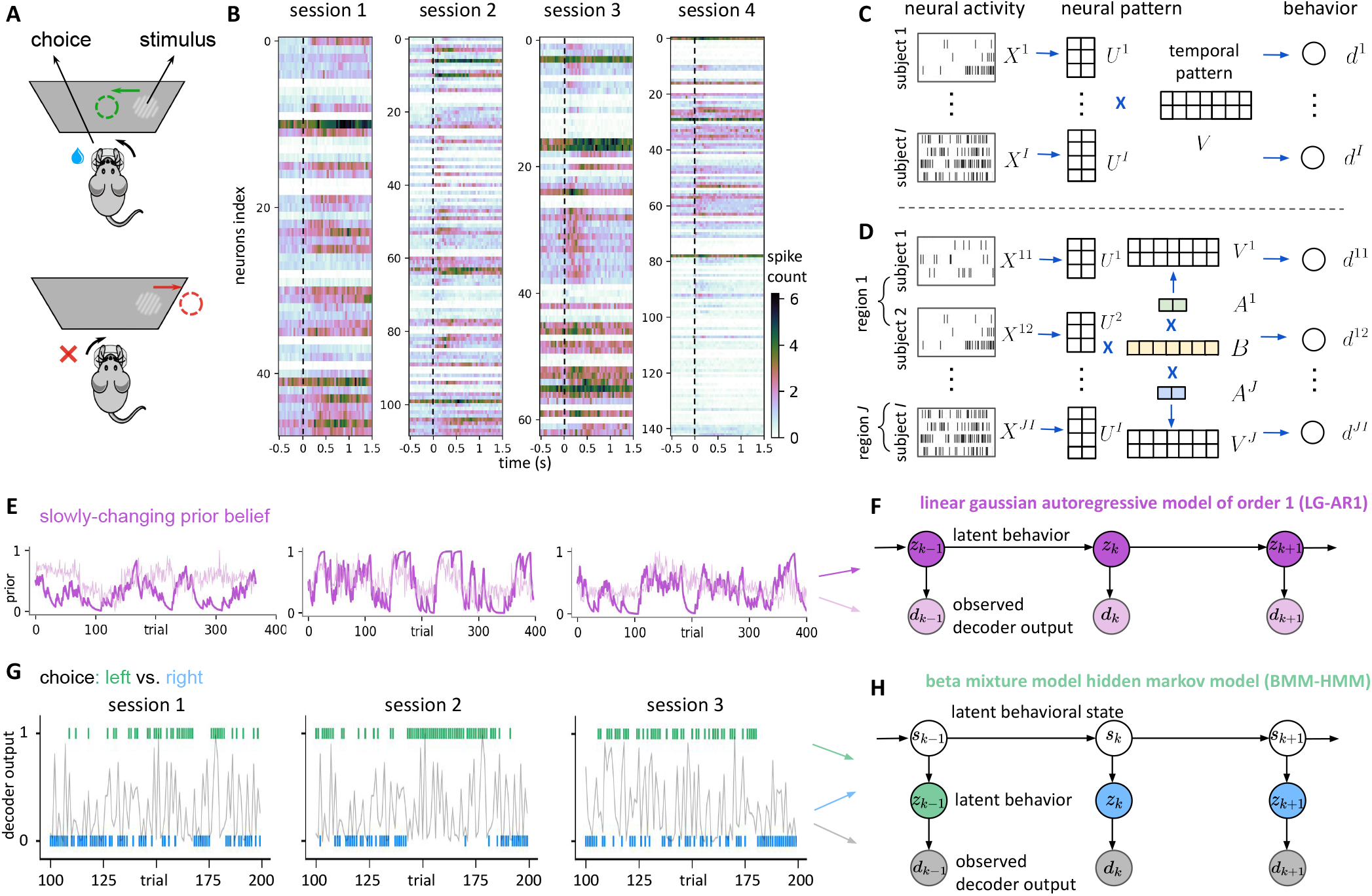
Schematic illustration of the neural and behavioral data-sharing models. **(A)** Schematic of the experiment where mice indicate the location of a visual stimulus by rotating a wheel. **(B)** Neural activity shows consistent activation following stimulus onset (dashed line at time t = 0s) across 4 selected sessions. Each spike train raster plot depicts the average spike count across all trials in a session. Each row in the plot represents the Peri-Stimulus Time Histogram (PSTH) of a single neuron. **(C)** Schematic of multi-session reduced-rank regression model. The blue symbol × denotes a multiplication operation. Neural activity from subject *i*, denoted *X*^*i*^, is first projected onto a low-dimensional subspace using a subject-specific neural basis *U*^*i*^, and then multiplied by a shared temporal basis *V* to produce the behavioral output *d*^*i*^. **(D)** Schematic of multi-region reduced-rank regression model. For each subject *i* and brain region *j*, neural activity is first projected onto a low-dimensional subspace using a subject-specific basis *U*^*i*^. This projection is then multiplied by a region-specific *V*^*j*^ to produce the behavioral output *d*^*ji*^. The temporal basis *V*^*j*^ is constructed by multiplying a region-specific weight matrix *A*^*j*^ with a shared global temporal basis *B*. **(E)** For slowly-changing prior belief *y*_*k*_ (dark purple), trial-to-trial correlations exist which single-trial decoders (light purple) neglect. Behavioral patterns are similar across sessions. **(F)** The LG-AR1 graphical model features latent behaviors *z*_*k*_ and observed single-trial decoder outputs *d*_*k*_, with colors corresponding to the examples in panel E. **(G)** For binary choice *y*_*k*_ (blue and green), trial-to-trial correlations exist, which single-trial decoders *d*_*k*_ (grey) fail to capture. Similar behavioral patterns also occur across sessions. **(H)** The BMM-HMM graphical model features latent behavioral states *s*_*k*_, latent behaviors *z*_*k*_, and observed single-trial decoder outputs *d*_*k*_, with colors corresponding to the examples in panel G.

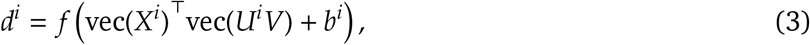

where 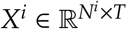 and *d*^*i*^ ∈ ℝ are the neural activity and predicted behavior from a single trial in session *i* with *N*^*i*^ neurons, corresponding to the terms *X* and *d* in Equation 2. As *V* is shared across sessions, a more robust estimation can be obtained since fewer parameters need to be learned from the same amount of data.

The multi-session reduced-rank regression model, by sharing a temporal basis across sessions covering diverse brain regions, assumes uniform neural activation patterns across regions. However, different brain regions may activate at different time steps within a trial due to functional differences (Kiebel et al., 2008; Scott et al., 2017; Zeraati et al., 2023). For example, sensory-related areas might activate earlier than cognition-related areas. To capture potential differences in temporal activation across brain regions while still enjoying the benefits of a low-rank model that combines information across multiple sessions, we propose a **multi-region reduced-rank regression model** (Figure 1D), decomposing the across-session temporal basis *V* into two low-rank matrices, allowing flexible temporal bases for different regions indexed by *j*:

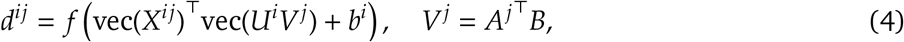

where 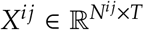 represents the neural activity from region *j* in session *i*, and *d*^*ij*^ ∈ ℝ is the behavior decoded from *X*^*ij*^. Intuitively, *A*^*j*^ ∈ ℝ^*L*×*R*^ captures regional differences, allowing varying timescales across brain regions. *B* ∈ ℝ^*L*×*T*^ represents shared similarities across regions, capturing major temporal variations associated with the behavior. In this context, *L* represents the rank of both the region-specific temporal basis set *A*^*j*^ and the global temporal basis set *B*. For *y*^*ij*^ ∈ ℝ, fitting a multi-session reduced-rank regression model (Equation 3) on *J* brain regions from *I* sessions learns *R × T* parameters for the temporal basis *V*. In contrast, fitting a multi-region reduced-rank regression model (Equation 4) on the same data slightly increases the temporal basis parameters to *L ×* (*J × R* + *T*). We select *L* and *R* via hyperparameter tuning, and empirically find that this procedure consistently outputs values of *L, R* < 10. This approach enables region-specific temporal bases to flexibly accommodate each brain area.

## 3 Formulation of the behavioral data-sharing model

In neuroscience experiments, animal behaviors often display trial-to-trial correlations. We can exploit these correlations to improve traditional single-trial decoders. For example, when neural activity from a single trial or session is insufficient for accurate decoding, incorporating information from neighboring trials or additional sessions can help improve decoder performance.

For traditional decoders, we use neural activity *X*_*k*_ in trial *k* to make predictions about the true behavior *y*_*k*_, and obtain a decoder estimate *d*_*k*_. The index *k* emphasizes that *X*_*k*_, *y*_*k*_ and *d*_*k*_ are single-trial quantities, with *X*_*k*_ corresponding to *X* in Equation 2 and *X*^*i*^ in Equation 3, and *d*_*k*_ corresponding to *d* in Equation 2 and *d*^*i*^ in Equation 3. (We focus on per-trial decoded scalar quantities *d*_*k*_ ∈ ℝ in this section, but this can be generalized.) Our goal is to improve *d*_*k*_ produced by the baseline decoder, which generates independent *d*_*k*_ for each trial. For example, when the baseline decoder is a traditional linear model, it captures correlations across neurons and time steps, but not across trials. Similarly, in the reduced-rank regression model, the neural basis *U* captures inter-neuronal dependencies, and the temporal basis *V* captures temporal structure within trials. However, neither model captures trial-to-trial dependencies.

Therefore we propose an approach to improve *d*_*k*_ by exploiting trial-to-trial correlations in *d*_*k*_ across all trials, and the statistical structure present in multiple sessions. Our method assumes that observations *d*_*k*_ are generated from latent variables *z*_*k*_ representing the unknown behavior, which follow a latent dynamic process. For continuous-valued behavior (e.g., an animal’s prior belief about stimulus side probability (Findling et al., 2023)), we model the transitions of *z*_*k*_ between trials using a first-order autoregressive process. Here, *z*_*k*_ in the current trial depends on *z*_*k*−1_ from the previous trial, while the continuous-valued *d*_*k*_ ∈ ℝ linearly depends on the latent *z*_*k*_ in the same trial. This is a **linear Gaussian autoregressive model of order 1 (LG-AR1)**. Given the sequence of decoder estimates 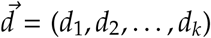, we can infer the latent variable *z*_*k*_ via standard Kalman smoothing forward-backward inference (Welch et al., 1995). This inferred *z*_*k*_ serves as an improved decoder estimate, potentially closer to the true behavior *y*_*k*_ than the single-trial estimate *d*_*k*_, by incorporating information from neighboring trials and other sessions. For the data generating mechanism, see Figure 1 E-F and STAR Methods, section “LG-AR1: Model details”.

While the LG-AR1 / Kalman smoother can provide improved estimates of continuous-valued *y*_*k*_ from noisy single-trial decoder estimates *d*_*k*_, this model is not applicable to binary-valued *y*_*k*_ ∈ {0, 1}, such as an animal’s choice in IBL’s experimental setup (IBL et al., 2022, 2023). In the IBL experiments, mice indicate the location of a visual stimulus by rotating a wheel. The stimulus appears randomly on either side with equal probability for the first 90 trials, then predominantly on one side (left or right) over blocks of subsequent trials. This setup creates a three-level data generating mechanism: (1) The animal forms an internal belief about the stimulus-generating behavioral state (*s*_*k*_); (2) Different choices (*z*_*k*_) are made based on the animal’s perceived state; (3) The decoder estimate *d*_*k*_ is generated depending on *z*_*k*_. This hierarchical structure requires a different modeling approach than LG-AR1.

For binary *y*_*k*_ ∈ {0, 1}, the output from single-trial decoder *d*_*k*_ ∈ [0, 1] represents the probability of *y*_*k*_ = 1. Our method assumes that *d*_*k*_ is generated from a mixture of beta distributions, with the mixture assignment dependent on the latent variable *z*_*k*_. When the single-trial decoder accurately predicts the behavior from neural signals, we expect well-separated beta mixture components. Specifically, *d*_*k*_ should be distributed close to 1 when *z*_*k*_ = 1 correctly predicts the true *y*_*k*_ = 1, and close to 0 when *z*_*k*_ = 0 correctly predicts the true *y*_*k*_ = 0. Conversely, if the decoder struggles due to insufficient neural information, the two beta distributions in the mixture become less distinguishable. We further assume that the latent variable *z*_*k*_ depends on latent behavioral states *s*_*k*_, whose transitions are governed by a hidden Markov model with *H* discrete hidden states. For instance, in the IBL binary choice task, at least three hidden states exist: random switching (stimulus appears randomly), left-biased, and right-biased (stimulus predominantly appears on one side). The likelihood of *z*_*k*_ being 0 or 1 varies with the latent state, defined by emission probabilities. We term this model the **beta mixture model hidden Markov model (BMM-HMM)**. Given the sequence of decoder estimates 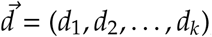, we infer both *s*_*k*_ and *z*_*k*_. The inferred *z*_*k*_ serves as an improved decoder estimate, potentially closer to the true behavior *y*_*k*_ than the original *d*_*k*_, by incorporating information from neighboring trials and other sessions. For the data generating mechanism, see Figure 1 G-H and STAR Methods, section “BMM-HMM: Model details”.

Single-session LG-AR1 and BMM-HMM models may output inaccurate parameter estimates when neural signals in the target session are insufficient. To address this, we propose multi-session versions of these models that utilize shared statistical structure across sessions to improve parameter estimation. Our multi-session approach learns empirical prior distributions of model parameters using observed behaviors from training sessions, and applies these priors to constrain model parameter updates during inference on the test session. This method, which borrows from empirical Bayes literature (Deely and Lindley, 1981; Robbins, 1992; Efron, 1996), pools data more effectively to constrain model parameters and improve characterization of underlying dynamics (Allenby and Rossi, 2006; Kruschke and Vanpaemel, 2015). For more information on prior distribution choices and implementation details, see STAR Methods sections: “BMM-HMM: Model details” and “LG-AR1: Model details.”

## 4 Results

We apply the proposed models to 433 sessions in the IBL datasets (IBL et al., 2022), covering 270 brain regions and 4 behavioral variables: choice, prior, wheel speed and whisker motion energy, which we describe in detail below. Although our main experiments focus on IBL data, the proposed methods are broadly applicable to settings where neural activity shows consistent temporal structure during the same behavioral tasks, and behaviors exhibit trial-to-trial correlations across sessions. To show this, we apply our models to the Allen Neuropixels dataset, which exhibits consistent temporal structure across repeated trials in a visual coding task. (Section 4.6). Furthermore, we demonstrate that our methods generalize to contexts without such structure, using IBL trial-unaligned data during spontaneous behaviors and a monkey random target reaching task without repeated trials.

In the IBL experiments, mice rotate a wheel to indicate the location of a visual stimulus, which is considered their *choice* (Figure 1A). For the first 90 trials, the stimulus appears randomly on either the left or right side of the screen with equal probability. In the subsequent trials, the stimulus appears predominantly on one side (either left or right) over blocks of trials (IBL et al., 2022, 2023). The mice are learning and adapting their behavior based on the changing probabilities in the experiment. This adaptive behavior allows us to estimate each mouse’s “prior belief” (*prior*) about the probability of where the stimulus appears per trial. The prior we consider is not the actual probability of stimulus occurrence. Instead, it represents an estimate of this probability for the current trial, based on the mouse’s behavior; see Findling et al. (2023) for details. *Wheel speed* and *whisker motion energy* near the whisker pad are also recorded. Whisker motion energy is extracted by computing the mean absolute difference between adjacent video frames in the whisker pad area (Laboratory et al., 2022), defined using a bounding box anchored between the nose tip and the eye. Choice and prior are static within a trial, while wheel speed and whisker motion energy are time-varying signals sampled at 60 Hz. For further details, refer to STAR Methods sections: “Data processing” and “Hyperparameter selection.”

### 4.1 Learning behaviorally relevant neural variations across sessions

The reduced-rank regression (RRR) model improves decoding by projecting neural activity onto a low-dimensional subspace that isolates behaviorally relevant variability. Unlike principal component analysis (PCA) (Abdi and Williams, 2010), which captures both task-relevant and irrelevant components (Cunningham and Yu, 2014; Sani et al., 2021), RRR emphasizes neural representations most predictive of behavior (Kobak et al., 2016). This supervised dimensionality reduction allows RRR to better disentangle behaviorally relevant neural dynamics compared to unsupervised methods like PCA or demixed PCA (Kobak et al., 2016); see STAR Methods, “Connection to PCA, CCA, demixed PCA, and sliced TCA”.

To illustrate this, we compare PCA and RRR projections of neural activity for decoding the binary choice variable. In Figure 2A, both PCA and RRR are applied to data from a single session, and the projections shown correspond to that session only. Different behavioral classes are more cleanly separated in the low-dimensional subspace of RRR than in PCA. Using K-means clustering and adjusted Rand index (ARI) (Hubert and Arabie, 1985; Steinley, 2004) to quantify class separability, we find that multi-session RRR produces more discriminative neural representations. This improvement arises from leveraging shared structure across sessions, which reduces overfitting and enhances generalizability. As shown in Figure 2 B-C, multi-session RRR achieves higher ARI and decoding accuracy than single-session RRR (*p* < 0.002 via bootstrap resampling; see STAR Methods for details), confirming the statistical significance of these gains.

**Figure 2:**
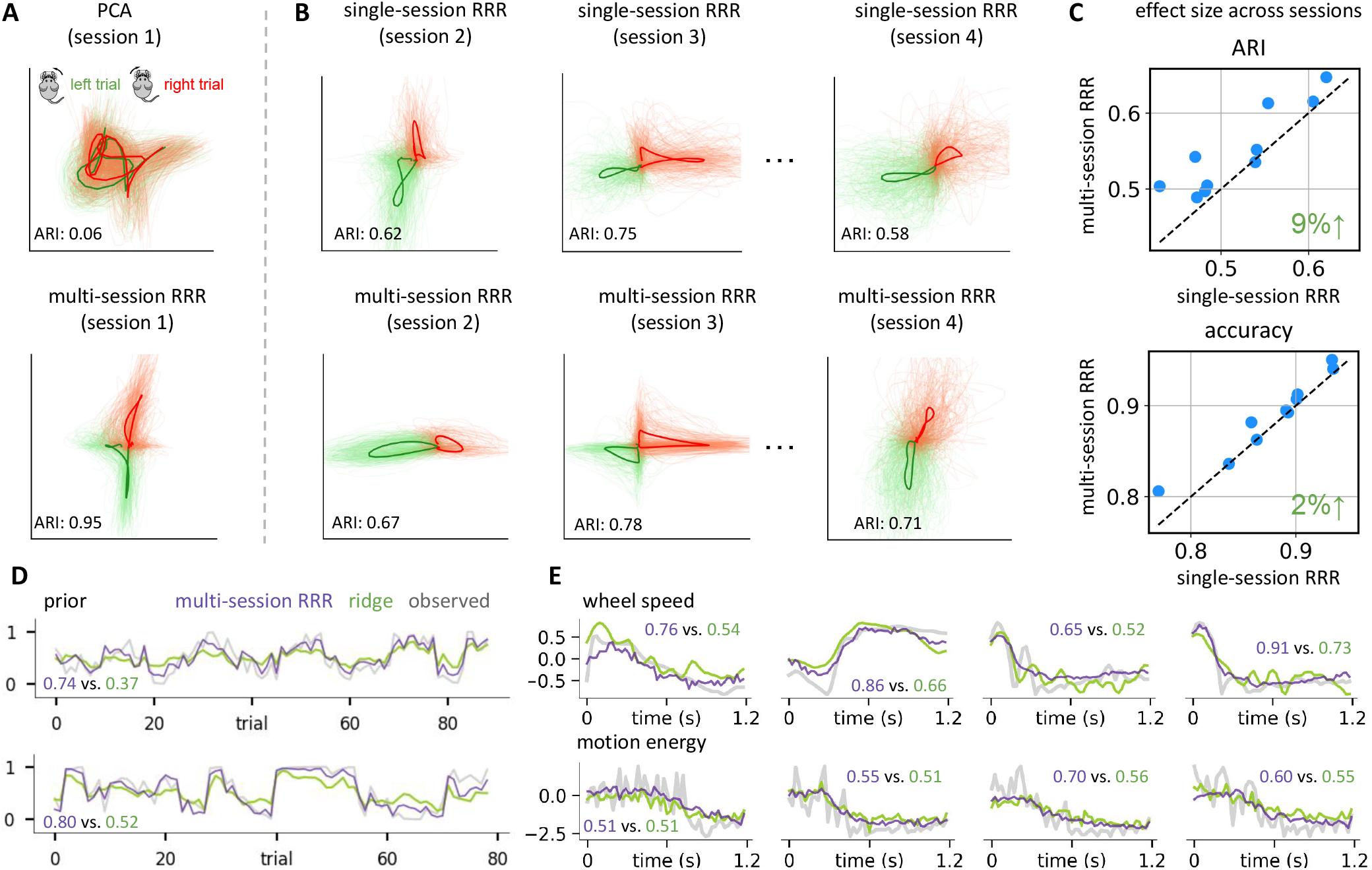
The reduced-rank regression model achieves strong decoding performance by learning behaviorally relevant neural variations through multi-session learning. **(A)** Projections of neural activity on the PCA subspace and the low-rank subspace identified by *U*^*i*^ from the multi-session reduced-rank regression model (RRR) are color-coded based on the binary behavioral variable. Light curves show single-trial projections from a single session, while dark curves represent trial-averaged projections. K-means clustering (2 clusters) is applied to the projections to separate left and right trials. Cluster similarity is assessed using the adjusted Rand index (ARI) (Hubert and Arabie, 1985; Steinley, 2004), where a higher score indicates better separation. Visualizations of the temporal basis *V* are depicted in Figure 7A. **(B)** Neural activity projections onto the low-rank subspace identified by the single-session and multi-session reduced-rank regression models, following the same color-coding convention as in panel A. K-means clustering is used to cluster the projections into left and right trials, and ARI measures cluster separation. **(C)** A scatterplot comparing ARI and decoding accuracy for the multi-session reduced-rank regression model versus the single-session variant across 10 sessions. Higher ARI values indicate better class separation. Each point represents an individual session. The green value in each subplot shows the relative improvement of the multi-session RRR over the single-session RRR. **(D)** Decoded prior from the multi-session reduced-rank regression model (purple) vs. ridge regression decoder (green), with Pearson’s correlation between the decoded and true prior shown as a numeric value for each example. The true prior (observed) is shown in grey. **(E)** Decoded motion energy and wheel speed traces from the reduced-rank regression model (purple) vs. ridge regression (green). The true behavior traces (observed) are shown in grey.

The reduced-rank regression model also improves decoding of continuous behaviors, such as prior, wheel speed, and whisker motion energy. Compared to ridge regression, RRR more accurately predicts these variables, as shown in Figure 2 D-E. We evaluate prediction quality in two ways: the ability to reconstruct trial-averaged behavior across stimulus conditions, and the ability to capture trial-specific variability beyond this average. Figure 3 A and I, B and J, and C and K show that RRR reliably reconstructs trial-averaged behavior across diverse conditions, while Figure 3 D and L, and E and M demonstrate that it also captures trial-specific variability after accounting for the trial average. Finally, analysis of residuals (Figure 3 F and N, G and O, and H and P) shows minimal systematic error for whisker motion energy, though slight underestimation of extreme wheel speeds. Despite this, the model accurately decodes continuous behaviors by capturing both trial-averaged structure and trial-specific variations.

**Figure 3:**
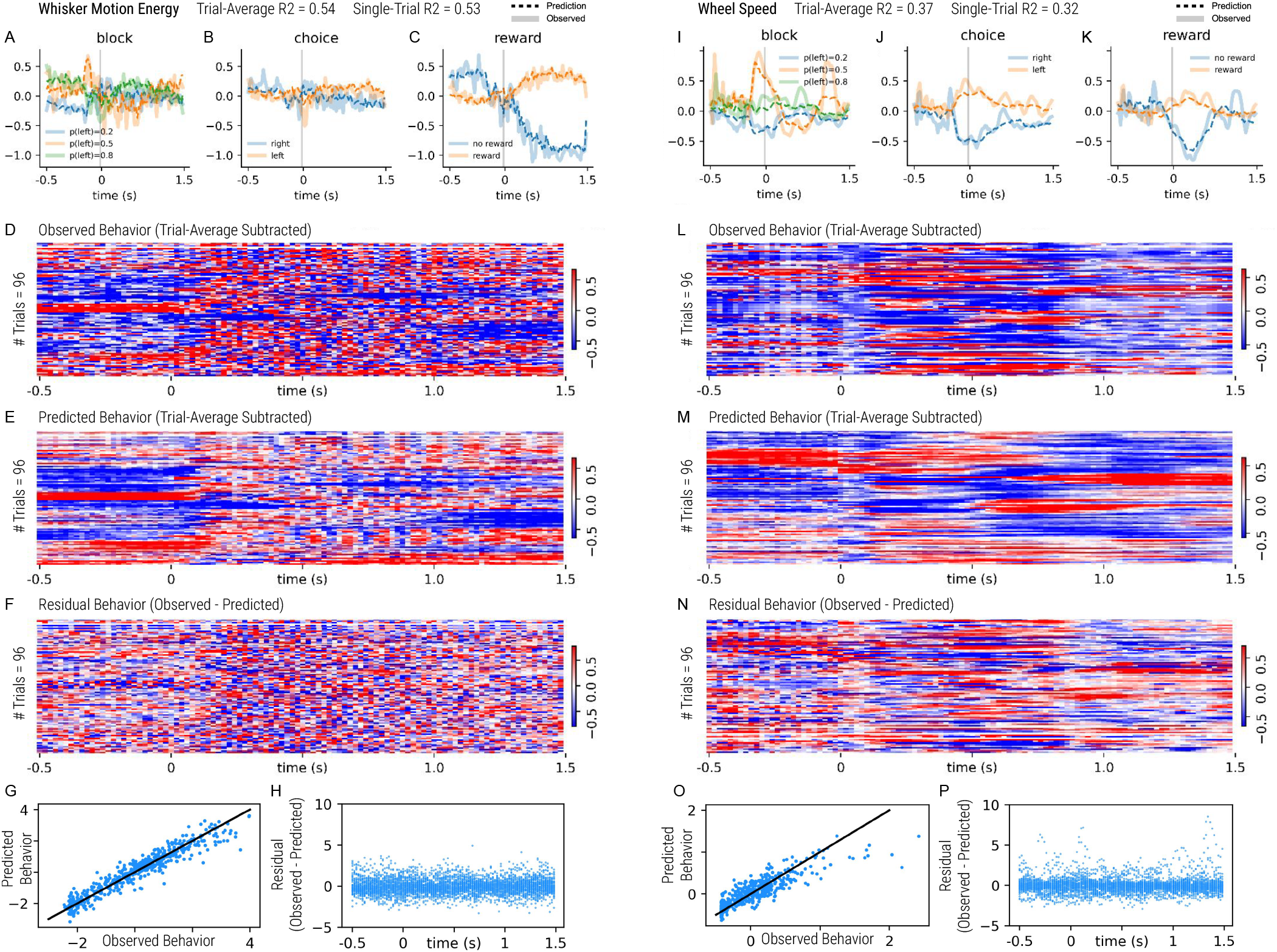
Qualitative evaluation of the reduced-rank regression model for decoding wheel speed and motion energy in a single IBL session. **(A, I)** Comparison of the reduced-rank regression model’s predicted behavior (dotted lines) with the observed ground truth behavior (solid lines) across different block conditions. For instance, blue lines indicate the average predicted (dotted) and observed (solid) behavior for trials with a block value of 0.2. The gray vertical line marks stimulus onset. **(B, J)** Average predicted and observed behavior across trials, grouped by choice condition (right versus left). **(C, K)** Similar comparison, but grouped by reward outcome (rewarded vs. unrewarded trials). **(D-F)** and **(L-N)** present data from 96 example trials in a session. Panel D shows the observed behavior, panel E displays the behavior predicted by the reduced-rank regression model, and panel F depicts the residuals (observed minus predicted behavior). In each panel, rows correspond to individual trials and columns to time steps within each trial. To highlight trial-to-trial variability, both observed and predicted behaviors are mean-centered by subtracting their respective trial averages. For visualization, all values (observed, predicted, and residual) are standardized. Spectral clustering is applied to the observed behavior to group trials with similar patterns. **(G, O)** Scatter plots showing the decoding performance of the reduced-rank regression model for whisker motion energy and wheel speed. Each point corresponds to a single trial and reflects the relationship between the observed (trial-averaged subtracted) and predicted behaviors (trial-averaged subtracted). For each trial, observed and predicted values are averaged over all time steps. A diagonal line is included to aid visual comparison between predicted and actual behavior. The predicted wheel speed tends to underestimate the actual wheel speed in many trials. **(H, P)** Distribution of residuals (observed minus predicted) over the time course of a trial. Residuals for whisker motion energy are centered around zero, indicating no systematic bias in the prediction errors. In contrast, residuals for wheel speed exhibit structured deviations from zero, suggesting the presence of some bias in the model’s predictions at high speeds.

### 4.2 Learning latent behavioral dynamics across trials

Next we turn to the behavioral data-sharing model. This model learns latent behavioral states 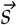 that infer the unknown behavior 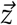 (Equation 21) given the neural activity *X*, leveraging the correlation between trials in the same state to improve single-session and single-trial decoder outputs 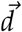. Figure 4A shows the latent state inference of a multi-session BMM-HMM applied to the IBL binary decision behavior (IBL et al., 2022). Recall that the stimulus probability switches between three discrete states: 1) a right (R) state (stimulus predominantly on the right), 2) a left (L) state (stimulus mostly on the left), and 3) a “middle” (M) state (stimulus randomly switching sides). Note that the three stimulus-generating states discussed here are different from the three decision-making states (“engaged,” “disengaged,” and “biased”) in Ashwood et al. (2022). The model accurately infers the occurrence of the three discrete states using only single-trial decoder outputs, without prior knowledge of the true choices or the timing of the stimulus probability block state changes. (Figure 4B). Note that we only use neural data from the decoded session to learn the model (the behavior in that session is unobserved). However, we do use observed behavior from other sessions to learn the multi-session model.

**Figure 4:**
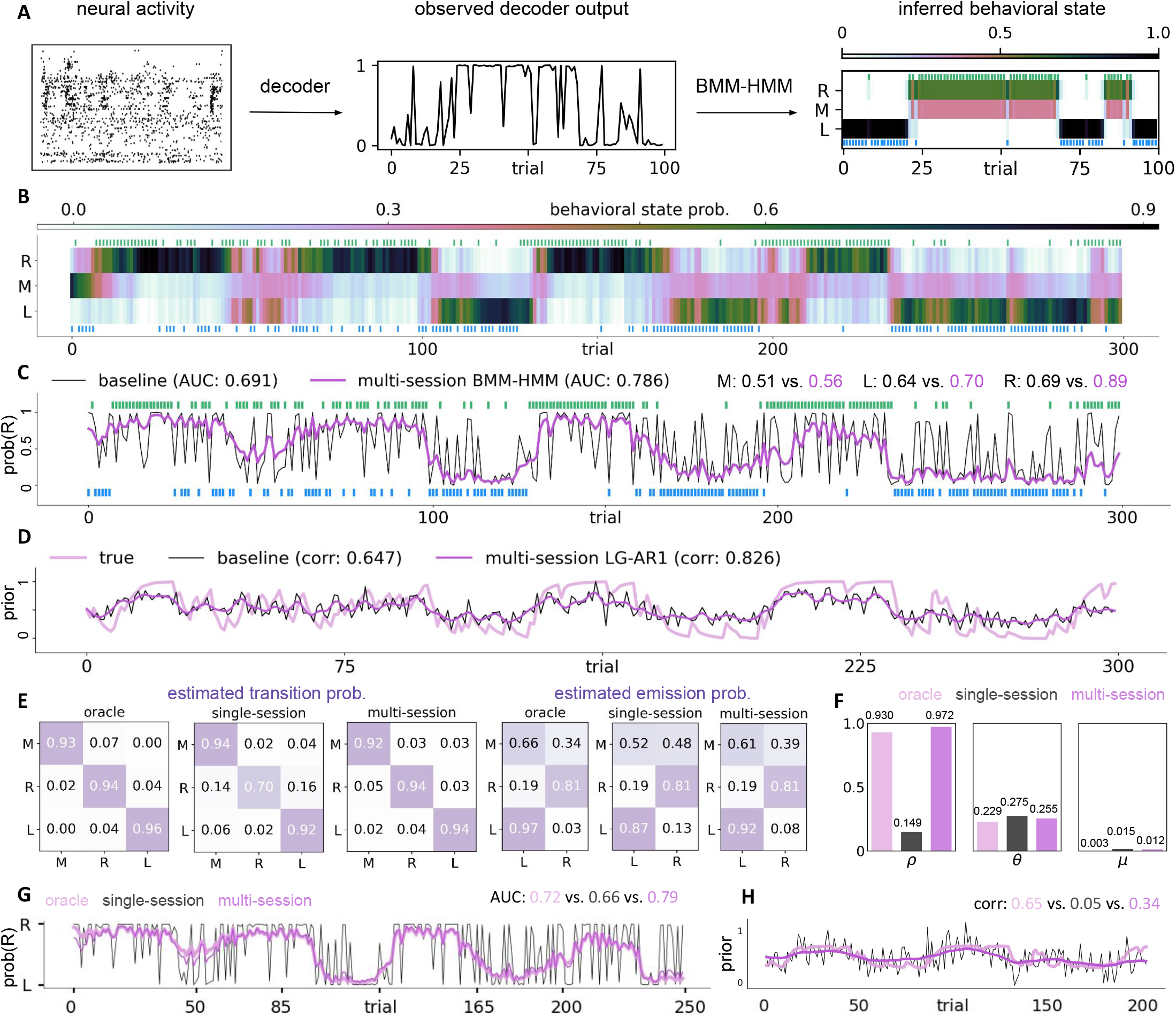
The behavioral data-sharing model improves single-trial decoding by inferring latent behavioral states from trial-to-trial correlations within individual sessions, and sharing behavioral information across sessions. **(A)** A schematic showing the BMM-HMM’s latent state inference from neural activity. A decoder is fitted to single-session, single-trial activity *X*_*k*_, yielding decoder output *d*_*k*_. The BMM-HMM is fitted to *d*_*k*_ to infer latent states *s*_*k*_, which alternate between left (L), right (R), and a random “middle” switching state (M), producing an improved decoder output. **(B)** The latent states *s*_*k*_ estimated from neural activity exhibit “block” structures, switching between states L, R, and M; these blocks mirror the true block probabilities in the IBL task but note that these states are learned, not pre-specified, and the state names in the plot are assigned post hoc. Color bar indicates state probabilities. Observed mouse choices are shown in green (right trials) and blue (left trials). **(C)** Improved decoder outputs 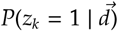 from the multi-trial and multi-session BMM-HMM (purple) overlaid on baseline single-trial and single-session decoder traces *d*_*k*_ (black), exploiting trial-to-trial correlations and achieving higher AUC. “Multi-session” refers to borrowing behavioral information from multiple training sessions to improve neural state estimates in the test session. *d*_*k*_ is observed and *z*_*k*_ is latent. We additionally show the decoder performance for each block type: random switching (M), left-biased (L), and right-biased (R). The decoding AUC of the baseline single-trial decoder is shown in black, while that of the BMM-HMM is shown in purple. **(D)** Improved decoder outputs 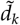 from the multi-trial and multi-session LG-AR1 (purple) superimposed on baseline single-trial and single-session decoder outputs *d*_*k*_, aligning more closely with the true prior (pink) and achieving higher Pearson’s correlation. **(E)** Estimated transition and emission probabilities from the oracle (pink), single-session (black), and multi-session (purple) BMM-HMM models. **(F)** Parameter estimates from the oracle, single-session, and multi-session LG-AR1 models. ρ is the first-order autocorrelation coefficient of the AR dynamical model, while θ and µ are the linear coefficient and bias of the Gaussian observation model (see STAR Methods, “LG-AR1: Model details”). **(G)** Decoded probabilities of choosing the right side from the oracle (pink), single-session (black), and multi-session (purple) BMM-HMM models. **(H)** Decoded priors from the oracle (pink), single-session (black), and multi-session (purple) LG-AR1 models.

Ideally, when the single-trial decoder accurately predicts behavior, the model can more precisely infer the states. In contrast, when the single-trial decoder makes errors, the model can compensate by borrowing decoder outputs from other trials (trial-to-trial correlation) and behavioral patterns from other sessions to refine its state estimation. Figure 4C visually compares the improved decoder outputs (Equation 21), from the multi-trial and multi-session BMM-HMM to the baseline single-trial and single-session decoder outputs. The single-trial and single-session decoder outputs exhibit considerable noise and frequent errors, while the multi-trial and multi-session outputs better follow the smooth “block” structure due to their knowledge of the latent states in the data. Quantitatively, the proposed model achieves a higher AUC (area under the ROC curve) than the baseline, highlighting the effectiveness of using trial-to-trial correlations and latent states to improve decoding. The decoder performance segregated by block type is also shown in Figure 4C. The decoding AUC of the baseline single-trial decoder is shown in black, while that of the BMM-HMM is shown in purple.

We apply similar ideas to improve the decoding of the continuous-valued *prior* using the LG-AR1 model. Recall that this prior signal represents a running estimate of the stimulus side probability (Findling et al., 2023), and tends to change smoothly over time. Similar to the BMM-HMM model, the LG-AR1 model infers the latent behavior (the true yet unobserved prior) by exploiting trial-to-trial correlations in the single-trial decoder outputs, borrowing behavioral information from other trials to correct estimates when decoding errors occur. Figure 4D visually compares the improved decoder outputs, 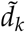 (Equation 51), from LG-AR1 to the baseline single-trial decoder outputs, *d*_*k*_. The single-trial decoder struggles to accurately predict the prior, as it doesn’t incorporate information from previous trials. In contrast, the LG-AR1 model, by considering trial-to-trial correlations, produces outputs that more closely align with the true prior, resulting in a higher Pearson’s correlation. Note that prior decoding is based on the full trial, so that the decoded signal may capture not only the prior variable but also correlated variables such as reward or visual stimulus.

We measure the impact of incorporating behavioral information from other sessions on the performance of the BMM-HMM and LG-AR1 models by exploring three model variants: a *single-session model*, a *multi-session model*, and an *oracle model* that uses true behaviors to learn parameters and improve decoder estimates (see STAR Methods for details). The oracle models assume that the true values of the latent behavioral variable *z*_*k*_ are known a priori. In this scenario, rather than inferring the latent behaviors, we directly substitute the ground truth observed behaviors *y*_*k*_ for *z*_*k*_, effectively treating *z*_*k*_ as a known quantity. However, the oracle models cannot simply use the observed *y*_*k*_ as the final improved decoder output, as this would result in a trivial decoding problem. Instead, these models must still generate a distinct output given the known *z*_*k*_ values and the learned model parameters. Thus the oracle model serves as an upper bound to assess the performance of single-session versus multi-session models. Figure 4E and F compare the estimated parameters of BMM-HMM and LG-AR1 from the three variants, showing that parameters estimated by the multi-session model align more closely with the oracle estimates than those from the single-session model. In addition, Figure 4G and H compare the outputs of the model variants, suggesting that predictions from the multi-session model are closer to the oracle model predictions than those from the single-session model. These findings underscore the importance of multi-session learning in improving both parameter estimation and decoding performance.

### 4.3 Benchmarking neural and behavioral data-sharing models with single- and multi-session baselines

To evaluate the impact of our proposed models on decoding performance, we perform a comprehensive benchmark encompassing both single-session and multi-session models. Single-session models include L2-regularized linear decoders, nonlinear multi-layer perceptron (MLP) decoders (Glaser et al., 2020), as well as single-session reduced-rank regression model, BMM-HMM, and LG-AR1. Multi-session models include the multi-session reduced-rank regression model, multi-region reduced-rank regression model, multi-session BMM-HMM and LG-AR1 models, along with a “combined” decoder, in which initial predictions are made using the multi-session reduced-rank regression model, and subsequently refined using either the multi-session BMM-HMM or LG-AR1 models. See STAR Methods for details on hyperparameter tuning and model architectures.

We decode choice, prior, wheel speed and whisker motion energy using neural activity from 5 selected brain regions in the reproducible electrophysiology (RE) datasets (IBL et al., 2022): the posterior thalamic nucleus (*PO*), lateral posterior nucleus (*LP*), dentate gyrus (*DG*), cornu ammonis (*CA1*), and anterior visual area of the visual cortex (*VISa*). These regions were chosen for their high neuron yield and anatomical diversity. To prevent ceiling effects, where decoding accuracy saturates when using all brain regions, we adopt a per-region decoding approach. This ensures more meaningful comparisons across models, as the selected regions individually contain less information than the full set of regions, resulting in more moderate and differentiable decoding performance.

For choice decoding, Figure 5A shows that the single-session reduced-rank regression model outperforms all other single-session baselines, including the nonlinear MLP. Despite hyperparameter tuning, the MLP may not have reached optimal performance, underscoring the advantage of our models, which have fewer parameters and thus allow for more thorough exploration of the model parameter space. The multi-session reduced-rank regression model (RRR (M)) provides further improvement over the single-session model, and the multi-region reduced-rank regression model (RRR (MR)) offers a slight additional gain. The “combined” decoder achieves the highest performance across most brain regions. While the BMM-HMM models perform better than the linear baselines, their improvement is not as substantial as that of the reduced-rank regression models.

**Figure 5:**
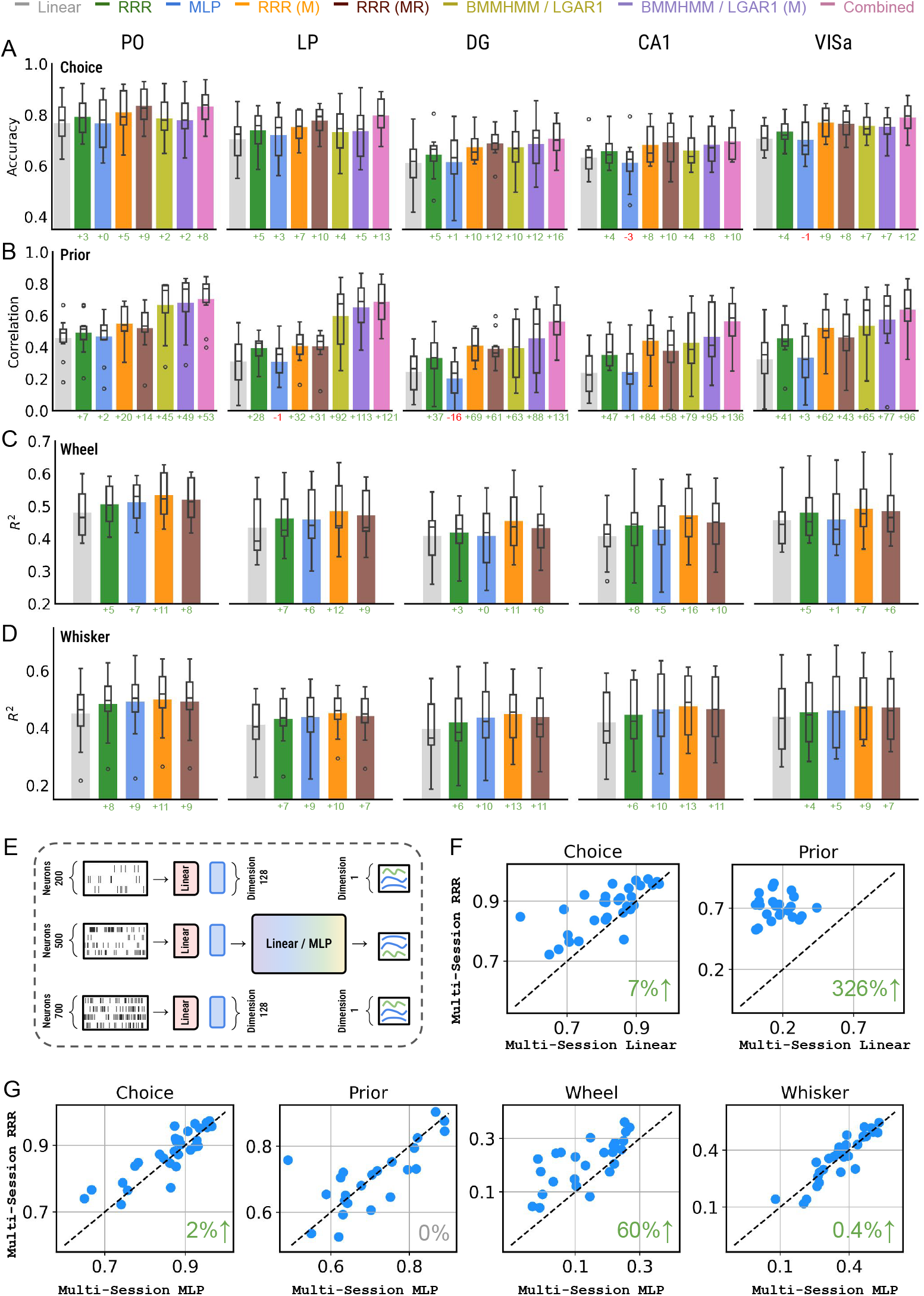
Quantitative comparison of the proposed neural and behavioral data-sharing models against single- and multi-session baselines. **(A-D)** Decoding performance for choice, prior, wheel speed and whisker motion energy using neural activity from 5 selected brain regions across 10 IBL sessions. The bar plots show the mean decoding metric, while the box plots show the median, quartiles, minimum, and maximum values of the metrics. For wheel speed and whisker motion energy, the *R*^2^ metric reflects how accurately single-trial behaviors are decoded. The multi-region reduced-rank regression model is denoted as “RRR (MR).” For the “combined” model, decoder estimates are first obtained using the multi-session reduced-rank regression model (RRR (M)), and then further refined with the multi-session BMM-HMM or LG-AR1 model (BMMHMM / LGAR1 (M)). The percentage improvement relative to the linear baseline is displayed as text at the bottom of the panel. The “combined” decoder is not benchmarked for wheel speed and whisker motion energy, as the multi-session BMM-HMM and LG-AR1 models have not been implemented for vector-valued continuous behaviors. **(E)** We use multi-session linear (full-rank) and MLP models as baselines for the multi-session reduced-rank regression models. To enable multi-session training across sessions with different numbers of neurons, we employ session-specific linear read-in matrices to project the neural activity from each session onto a latent space with the same dimension before feeding it into the full-rank linear or MLP model backbone. **(F-G)** A scatterplot comparing the multi-session reduced-rank regression model to the multi-session linear and MLP models, with all models trained on 30 sessions. Each dot represents an individual session. The green (gray) value in the bottom right of each subplot indicates the relative improvement of the multi-session reduced-rank regression model over the corresponding multi-session baseline. Decoding metrics for wheel speed and whisker motion energy are not displayed for the multi-session linear model due to its poor performance.

For prior decoding, the LG-AR1 models show improvements that are comparable to or greater than those achieved by the reduced-rank regression models (Figure 5B). The reduced-rank regression model outperforms other single-session baselines but falls short of the multi-session reduced-rank regression model. The multi-region reduced-rank regression model performs slightly below the multi-session variant. The “combined” decoder, which integrates the multi-session reduced-rank regression and LG-AR1 models, achieves the highest decoding performance across all models. For wheel speed and whisker motion energy decoding, Figures 5C and D show that the single-session reduced-rank regression model outperforms the linear baselines and performs comparably to the MLP decoders. The multi-session reduced-rank regression model achieves the best overall performance, while the multi-region model shows slightly lower performance than the multi-session variant.

Above we observe that multi-session models consistently outperform the single-session models. This raises a natural question: Is the improved decoding performance due to the structure of the reduced-rank regression model—specifically, its ability to capture shared temporal patterns across sessions—or simply a result of training on more data? To disentangle these factors, we compare the multi-session reduced-rank regression model with two alternative multi-session baselines: full-rank linear and MLP decoders. To support multi-session training across sessions with varying neuron counts, we utilize session-specific linear read-in layers to project neural activity from each session into a latent space with the same dimension, before feeding the latent representation into the full-rank linear or MLP model (Figure 5E). Figure 5F shows that the multi-session reduced-rank regression model outperforms the multi-session linear model in decoding both choice and prior. Given the poor performance of the multi-session linear baseline, we exclude its results for wheel speed and whisker motion energy from the analysis. In Figure 5G, the multi-session reduced-rank regression model outperforms the multi-session MLP in decoding choice, wheel speed, and whisker motion energy, while achieving comparable performance in prior decoding. These results indicate that the performance gains of the multi-session reduced-rank regression model stem not only from multi-session training, but also from its structured model design.

### 4.4 Identifying important neurons for decoding

The reduced-rank regression model not only improves decoding outcomes but also offers intrinsic interpretability. In this section, we show that the neural basis *U* quantifies individual neurons’ contribution to behavior decoding (see Equation 12 for theoretical justification). We validate this claim through a “neuron pruning” experiment, where the magnitude of *U*’s first rank indicates neuron importance, with larger values indicating higher importance. Starting with all neurons, we iteratively remove 5% of neurons from each session. After each removal, we fit a L2-regularized logistic regression to the remaining neurons’ activities and track the decrease in decoding accuracy measured by AUC. We compare three removal strategies: removing the least important neurons first, removing the most important neurons first, and removing randomly selected neurons. Figure 6 A-E show that removing the least important neurons first minimally impacts decoding performance, while removing the most important ones leads to a faster decline in choice decoding accuracy than random removal. Moreover, accurate decoding can be achieved with only a small proportion of the important neurons (green curves in Figure 6 A-E). Figure 6 F-J show the choice-conditioned, trial-averaged activity of the most and least important neurons identified based on the reduced-rank regression model’s *U* values from example sessions in each region. The most important neurons exhibit choice-selective firing patterns, while the least important neurons show similar activity in left and right trials, indicating limited task responsiveness.

**Figure 6:**
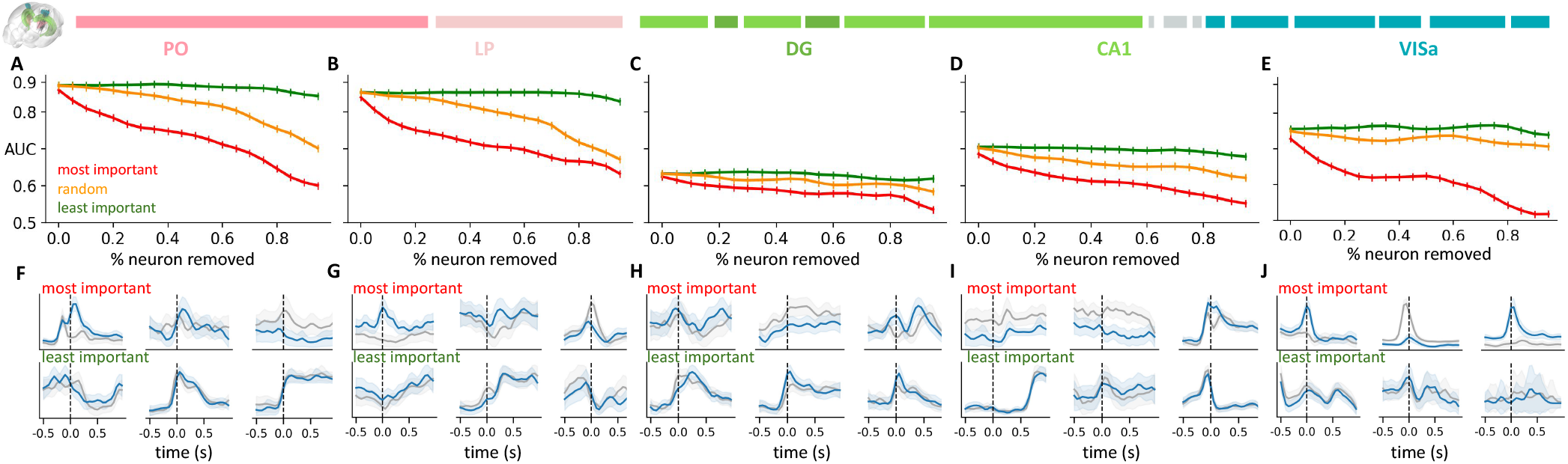
Reduced-rank regression models identify important neurons for decoding choice in brain regions including PO, LP, DG, CA1, and VISa. **(A-E)** Region-specific performance degradation from the “neuron pruning” experiment using three neuron removal strategies. Decoding accuracy is quantified by AUC and averaged across 10 sessions from each region. **(F-J)** Trial-averaged neural activities conditioned on choice for the most and least important choice-decoding neurons from example sessions in each brain region. Blue and black solid curves show the mean spiking patterns for left and right trials, respectively, with light-colored ribbons indicating one standard deviation. Stimulus onset is indicated by a vertical dashed line.

### 4.5 Mapping behaviorally-relevant timescales across the brain

Prior studies have shown that functionally distinct brain regions exhibit varying intrinsic timescales (Kiebel et al., 2008; Scott et al., 2017; Zeraati et al., 2023), with motor and sensory areas operating on faster timescales than cognition-related regions. However, a comprehensive examination of how these temporal dynamics relate to specific behaviors remains largely unexplored. To address this, we fit the multi-region reduced-rank regression model on 433 sessions across 270 brain areas to decode choice and prior, and represent the first rank of the region-specific temporal basis *V*^*j*^ as the timescale of each brain region. Figure 7A shows distinct activation timescales for different brain regions in decoding choice, including the gigantocellular reticular nucleus (GRN), motor cortex (MOp), nucleus accumbens (ACB), amygdala complex (CEA), CA1 region in the hippocampus, basomedial amygdala (BMA), and visual cortex (VISa). The peak activation time (*peak*) corresponds to the highest point of a curve. The activation duration (*width*) is defined as the interval spanning points on either side of the peak where the curve covers 90% of the peak height. Although peak activation occurs at similar times following stimulus onset, ACB and BMA show more prolonged activation compared to other regions.

**Figure 7:**
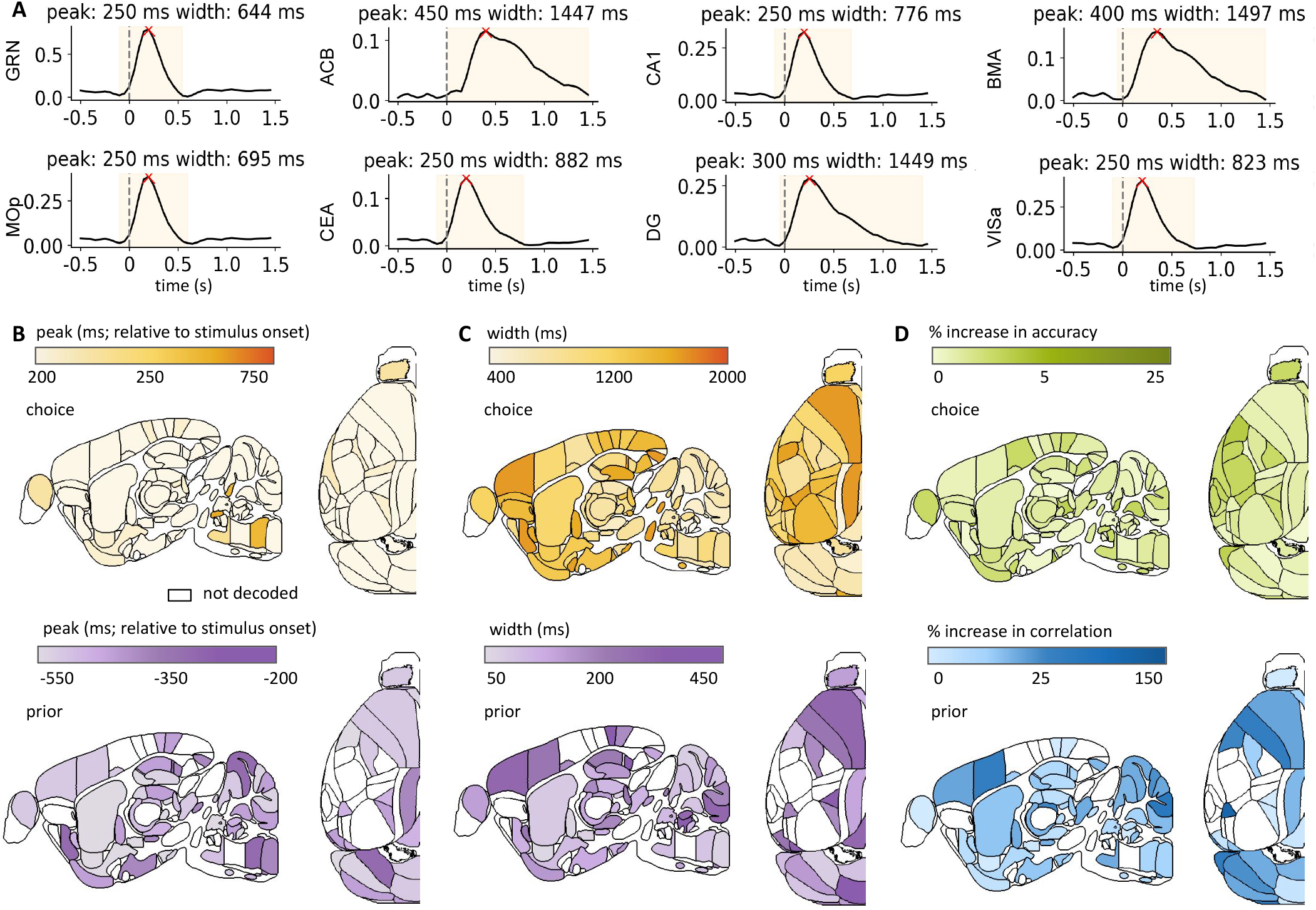
Mapping behaviorally relevant timescales and decoding quality improvement across the brain. **(A)** The first rank of each brain region’s temporal basis *V*^*j*^ in the multi-region reduced-rank regression model (Equation 4) is shown. Stimulus onset is indicated by a dashed line, peak activation time (“peak”) by a red cross, and activation duration (“width”) by a yellow segment. “Peak” corresponds to the highest point of a curve. “Width” is defined as the interval spanning points on either side of the peak where the curve covers 90% of the peak height. **(B)** Brain-wide map of relative peak activation time w.r.t. stimulus onset. **(C)** Brain-wide map of activation duration (width). Colors distinguish choice (yellow) from prior (purple); intensity represents peak time and duration magnitude. White regions indicate non-decoded areas. **(D)** Region-specific improvement in choice decoding accuracy and the correlation between the real and predicted prior. The multi-region reduced-rank regression model’s improvement is compared to the baseline L2-regularized linear decoder. Color intensity represents the magnitude of improvement.

We use the peak activation time and duration of each area (Figure 7A) to compare behaviorally relevant timescales across brain regions. Figure 7A shows that for the choice decoding task, most brain regions exhibit peak activation within 1.5 seconds of stimulus onset. This timing aligns closely with the “reaction time,” defined as the interval between stimulus onset and the initial movement (Figure 1c of IBL et al. (2023)). For the choice decoding task (visual decision-making), Figure 7B (first row) shows most regions have similar peak activation times, except the olfactory bulb and cerebellum, which may have delayed activation upon receiving the water reward. Figure 7C (first row) shows that activation durations vary, with hindbrain areas having shorter durations than forebrain and midbrain regions. For the prior decoding task (learning from past experiences), Figure 7B (bottom row) shows the cerebral cortex has earlier activation, while regions in the cerebellum have delayed activation. Figure 7C (bottom row) shows the cerebral cortex and thalamus have longer activation durations than other areas. White areas denote brain regions not decoded due to the absence of corresponding behavioral data (choice or prior) in sessions containing these regions.

In addition to showing when each brain region is activated by a behavioral variable and the duration of its activation in Figure 7 B and C, we analyze the amount of decodable behavior information from the neural activity in each region. While IBL et al. (2023) creates a brain-wide map of decoding accuracy for selected behavior tasks, they only use L2-regularized linear decoders. In Figure 7D, we show that the multi-region reduced-rank regression model, a more constrained and interpretable linear decoder trained with more data, improves choice and prior decoding across most brain regions compared to the linear decoder baseline used in IBL et al. (2023). This suggests that regularized linear decoders may not fully capture all decision-making task information in each region, potentially influencing the interpretation of results derived from these decoders.

In STAR Methods, section “Assessing statistical significance of decoding improvements via null distribution,” we confirm that multi-region reduced-rank regression model improves the amount of information decoded from each region relative to the baseline linear decoder. To control for potential spurious correlations (Harris, 2020), we generate null distributions using “imposter sessions” following the approach of IBL et al. (2023). Analysis of representative brain regions (PO, LP, DG, CA1, and VISa) in Figure S4 implies that while absolute decoding improvement varies slightly between original and adjusted scores, the relative ranking of regional improvements remains largely consistent.

### 4.6 Generalization across data structures, species, and behavior tasks

In the previous sections, we focus on IBL trial-aligned Neuropixels recordings of mice performing visual decision-making tasks. To demonstrate the generalizability of our models across diverse data structures, species, and behavior tasks, we extend our methods to three additional datasets: (1) IBL trial-unaligned data, which include neural activity during both structured, task-related engagement and spontaneous behaviors; (2) the Allen Institute Neuropixels visual coding dataset, which captures neural activity in mice exposed to visual stimuli different from those used in IBL tasks (de Vries et al., 2023); and (3) a primate random target task, involving self-paced, sequential reaching behavior without repeated trials (Pei et al., 2021).

In the prior experiments, we segment recordings into 2-second trials aligned to explicit time-based cues, such as stimulus onset or first movement onset. To demonstrate that our methods are not limited to data with well-defined trial structures, we apply them to IBL trial-unaligned data, which includes 2-second snippets sampled from both within and outside of experimental trials (Figure 8A). These snippets capture freely behaving periods, without alignment to specific task events. We decode two continuously recorded behavioral variables: wheel speed and whisker motion energy. As baselines, we compare ridge regression (linear baseline), a single-session reduced-rank regression model, a single-session MLP model, and a multi-session reduced-rank regression model. As shown in Figure 8B, the single-session reduced-rank regression model outperforms the linear baseline and achieves performance comparable to the nonlinear MLP. The multi-session reduced-rank regression model provides a modest but consistent improvement over its single-session version. Figure 8C displays the predicted behaviors from each model, showing that the multi-session reduced-rank regression model produces predictions that more closely follow the ground truth, indicating improved decoding accuracy.

**Figure 8:**
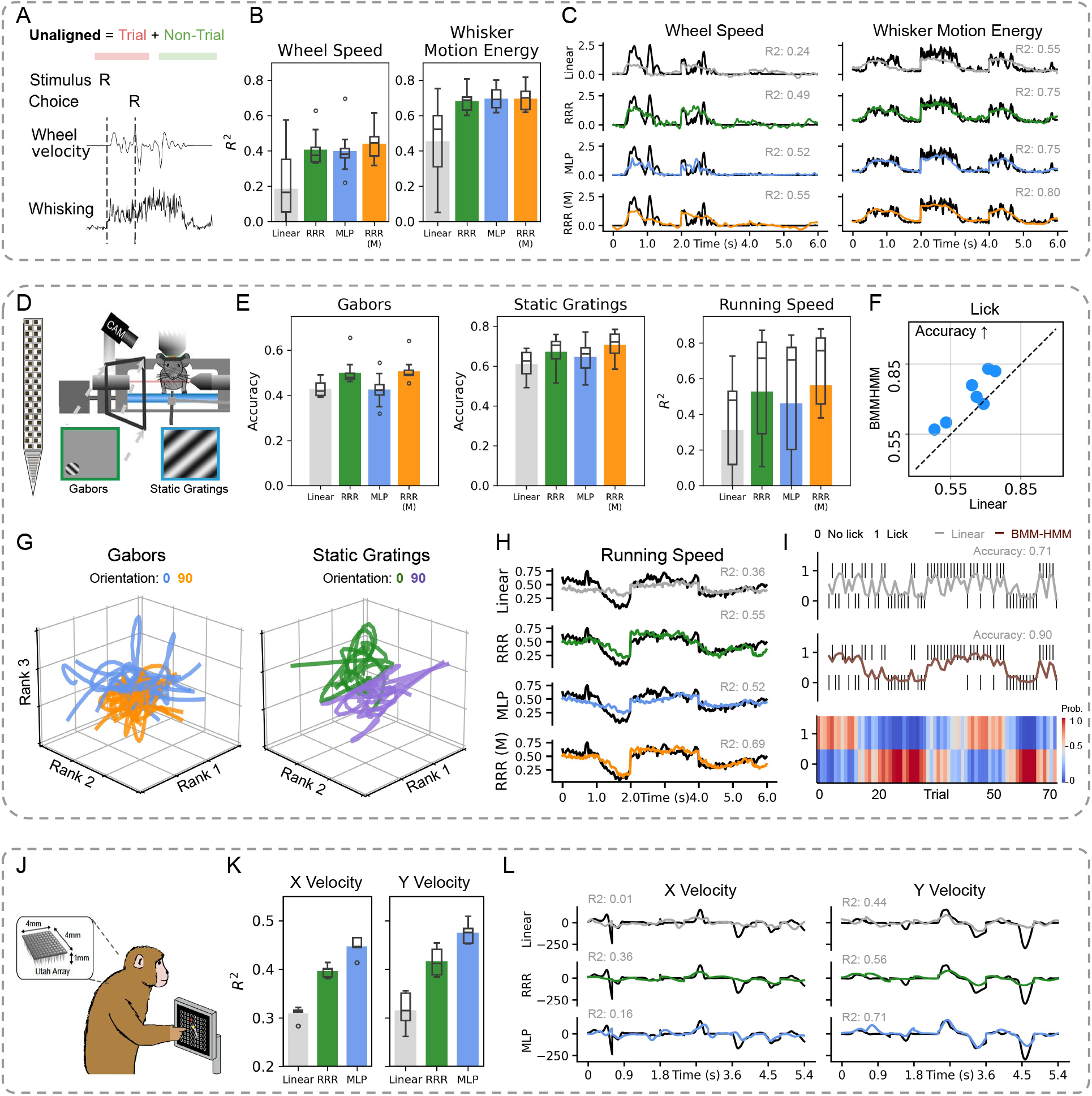
Generalization across data structures, species, and behavior tasks. **(A)** The IBL trial-unaligned dataset includes recording snippets sampled from both within and outside of structured experimental trials. **(B)** We compare the performance of single-session (*RRR*) and multi-session reduced-rank regression models (*RRR (M)*) to linear and MLP baselines across 10 IBL sessions. The *RRR (M)* model is trained using data from 50 IBL sessions. Bar plots represent the mean decoding metric, while box plots indicate the median, interquartile range, and full range (min to max). **(C)** Predicted wheel speed and whisker motion energy from an example session are compared to ground truth. Predictions from the multi-session model (*RRR (M)*) align more closely with the actual behavior. **(D)** The Allen Institute visual coding dataset includes Neuropixels recordings from mice exposed to different visual stimuli. **(E)** We evaluate decoding performance across 10 Allen sessions by benchmarking single-session and multi-session reduced-rank regression models against baseline models. The multi-session reduced-rank regression model (RRR (M)) is trained on data from 58 Allen sessions. Mean decoding metric is shown with bar plots, while box plots display the distribution, including median, quartiles, min and max. **(F)** Comparison of the BMM-HMM model and a linear baseline in predicting behavioral responses (licks) to visual stimulus changes across seven example sessions. Each dot represents the accuracy of a single session. **(G)** Neural activity from each trial, projected onto the latent space of the reduced-rank regression model, is color-coded by stimulus orientation for both Gabors and static gratings. For clarity, only two orientations are shown. **(H)** Predicted running speed from an example session is plotted alongside the ground truth, with the multi-session model (RRR (M)) capturing the actual behavior more accurately. **(I)** Side-by-side comparison of predicted and ground truth licks from the BMM-HMM model and a linear baseline in an example session. The corresponding inferred latent states are shown as a heatmap, color-coded by the probability of each state. **(J)** Neural activity is recorded using Utah arrays as the monkey engages in a random target reaching task. **(K)** Decoding performance for finger velocity along the X and Y axes is evaluated using linear, MLP, and reduced-rank regression models within a single session via 5-fold cross-validation. Bar plots show the mean decoding *R*^2^, while box plots display the distribution of *R*^2^ across folds, including the median, quartiles, min and max. **(L)** Finger velocity predictions along the X and Y axes are compared to the true recorded velocities.

To evaluate the generalizability of our models across different behavior tasks, we apply them to the Allen Institute visual coding dataset, which includes Neuropixels recordings from the visual cortex and thalamus of mice exposed to a broad range of visual stimuli, including Gabors and static gratings (Figure 8D). During experiments, mice are presented with Gabors at three orientations and static gratings at six orientations, making the decoding task a multi-class classification problem. Additionally, we decode running speed, a continuous, trial-unaligned behavioral variable spanning both stimulus presentations and free-behavior periods. We compare the performance of several baseline models across all behavior variables: an L2-regularized linear model, a single-session reduced-rank regression model, a single-session MLP, and a multi-session reduced-rank regression model. As shown in Figure 8E, the single-session reduced-rank regression model consistently outperforms the linear and MLP baselines, and the multi-session version slightly improves its performance. In Figure 8G, we visualize the low-dimensional neural projections from the reduced-rank regression model, revealing latent spaces that clearly separate stimulus conditions, indicating that the model captures behaviorally relevant neural variations. Figure 8H compares predicted and ground-truth running speed, showing that both the single-session and multi-session reduced-rank regression models decode this trial-unaligned behavior with high fidelity.

The BMM-HMM model infers latent internal states underlying observed behavioral outcomes, such as when a mouse perceives a stimulus and responds with a choice. Since Gabors and static gratings are stimuli, not behavioral responses, the BMM-HMM is less applicable to decoding these variables. To showcase BMM-HMM, we focus on a different “choice”-related variable: *licks*, which are behavioral responses to changes in visual stimuli. When the type of visual stimulus changes, the mouse is expected to detect this change and respond with a lick, which is rewarded if correct. However, mice often produce consecutive licks to maximize their probability of receiving a reward (de Vries et al., 2023). Figure 8I shows that BMM-HMM uncovers distinct “engaged” and “disengaged” states in a session. In the “engaged” state, the mouse licks consistently, whereas in the “disengaged” state, licking is infrequent. By inferring internal states that capture trial-to-trial correlations, BMM-HMM improves decoding performance over the linear baseline (Figure 8F). See STAR Methods “Data processing” and Figure S6 for data preprocessing details and additional visual examples.

In addition to the Allen dataset, we also apply our method to a monkey random target task (Pei et al., 2021) to demonstrate its generalization across recording hardware and species. In this self-paced, sequential reaching task, the monkey moves between random elements of a grid (Figure 8J). The dataset consists of single-session neural spiking activity recorded from the primary motor cortex using Utah arrays, along with simultaneously recorded finger positions. We decode finger velocity and compare the performance of a single-session reduced-rank regression model to two baselines: ridge regression and a single-session MLP model, using 5-fold cross-validation. As shown in Figure 8K, the reduced-rank regression model outperforms the linear ridge regression, though its performance is below that of the nonlinear MLP. Figure 8L shows predicted versus ground truth finger velocity along the X and Y axes for each model. Although the reduced-rank regression model does not outperform the MLP in this motor decoding task, it offers a substantial improvement over the standard linear decoder.

## 5 Discussion

We propose multi-session reduced-rank regression and state-space models to share neural and behavioral structures across sessions to improve neural decoding. When applied to a large collection of sessions spanning multiple brain regions, our models improve decoding performance on several behavior variables. In addition to being performant, our approach is interpretable: it highlights important neurons for decoding, identifies behaviorally relevant timescales in different brain areas, and infers latent behavioral states from neural activity. While our reduced-rank framework can be extended with nonlinear models, our focus in this work is on developing an efficient and interpretable linear decoder to replace traditional full-rank linear regression, while also enabling multi-session training. The reduced-rank structure provides clear interpretability through the neural and temporal bases *U* and *V*, and it often matches or exceeds the performance of simple nonlinear MLPs, especially when computational resources for hyperparameter tuning are limited (see STAR Methods for computation time comparison).

Several existing methods relate to our neural data-sharing model (Mante et al., 2013; Aoi and Pillow, 2018; Semedo et al., 2019, 2020; Syeda et al., 2023; Stringer et al., 2019). Gallego et al. (2018) uses canonical correlation analysis (CCA) to align latent dynamics across sessions, while we substitute the unsupervised CCA with the proposed reduced-rank regression model using a supervised decoding loss. CCA maximizes neural-behavioral correlation, but our model minimizes the normalized mean squared error between the real and predicted behavior. Demixed PCA (Kobak et al., 2016) isolates neural activity variations related to different conditions, maximizing neural-behavioral correlations and prioritizing neural variability for reconstruction. In contrast, our model emphasizes behavioral variation for accurate decoding. While the proposed model has a decoding target (behavior), demixed PCA can be viewed as a form of reduced-rank regression with an encoding target (neural activity). The preferential subspace identification (PSID) (Sani et al., 2021) also extracts low-dimensional, behaviorally relevant neural dynamics but rely on more complex state-space models. The proposed reduced-rank regression model is a latent variable model without constraints on neural dynamics. Additionally, the proposed model resembles methods in the tensor component analysis (Williams et al., 2018; Pellegrino et al., 2024) and tensor regression literature (Guo et al., 2011; Zhou et al., 2013). In particular, the factorization of the full-rank weight matrix into the low-rank matrices *U* and *V* can be viewed as a special case of low-rank tensor decomposition applied to a two-mode regression tensor. Although our approach is formulated in a matrix setting, it shares the same underlying motivation to reduce overfitting and improve generalization by constraining the model through low-rank structure.

Previous work has applied hidden Markov models to infer latent behavioral states from movement (Whoriskey et al., 2017; Wang, 2019) or decision-making data. For example, Ashwood et al. (2022) models mouse decision-making using HMMs with generalized linear model (GLM) observations, allowing behavioral states to persist across trials and depend on the stimulus and other covariates. Unlike these methods that infer HMM states only from the behaviors, we also use neural data. While Abeles et al. (1995); Kemere et al. (2008); Danóczy and Hahnloser (2005); Rainer and Miller (2000); Radons et al. (1994) apply HMMs to understand how different neural states generate the observed neural activities, we learn HMM states that generate the observed decoder estimates, which rely on both neural activity and behavior. Another related approach is that of Batty et al. (2019), which uses a Bayesian decoder to infer both continuous and discrete states from animal behavior video data, and then combine those with a behavior-based autoregressive HMM to smooth the original predictions. While we focus on showing the improvement from state-space models that exploit trial-to-trial correlations in IBL data, the reliance on task/behavior history is a common feature in many neuroscience experiments and datasets. For example, beyond fitting a GLM-HMM to capture consistent behavioral states across trials, Ashwood et al. (2022) also applied their method to a non-IBL mouse dataset and a human dataset focused on motion coherence discrimination. Additionally, animal behavior is inherently influenced by past experiences (Foucault et al., 2024; Sainburg et al., 2025). In these contexts, incorporating task and behavior history is valuable not only for decoding but also for studying learning and memory mechanisms.

Technological advances now enable the simultaneous collection of multiple data modalities, such as local field potentials and calcium imaging, during neuroscience experiments. Moreover, reduced-rank regression models have applications beyond neural decoding, including neural encoding (predicting neural activity from behavior) (Kobak et al., 2016; Steinmetz et al., 2019; Posani et al., 2025) and inter-region activity prediction (reconstructing activity in one brain region using data from another) (Semedo et al., 2022). Therefore, important future directions include incorporating more data modalities into the model and adapting the model to perform additional tasks. Another important future direction is relaxing the assumption of shared temporal bases across sessions in the reduced-rank regression model. While our current approach assumes a shared temporal structure across sessions to enhance decoding performance, this assumption may not always hold, particularly in scenarios where animals are at different learning stages or display varying performance levels between sessions. For multi-session state-space models, exploring nonlinear time series models and dynamical systems (Hochreiter and Schmidhuber, 1997; Chen et al., 2018; Rubanova et al., 2019) can facilitate modeling more complex latent behavioral dynamics. Finally, all of our models are compatible with the density-based decoding approach from Zhang et al. (2024a), allowing decoding from unsorted spike features rather than spike-sorted data; we expect that combining these approaches would lead to further accuracy improvements.

## Acknowledgments

We thank Matt Whiteway for many useful discussions, Peter Latham and Jonathan Pillow for helpful comments on the manuscript, and Chris Langfield for code review. This work is supported by grants from the Wellcome Trust (209558 and 216324), National Institutes of Health (1U19NS123716 and K99MH128772), the Simons Foundation, and the National Science Foundation and DoD OUSD (R&E) under Cooperative Agreement PHY-2229929 (The NSF AI Institute for Artificial and Natural Intelligence).

## STAR Methods

### Method details

#### Mathematical notation

Throughout the main text, STAR Methods section, and supplemental information, we use the mathematical notation defined in Table 1.

**Table 1:**
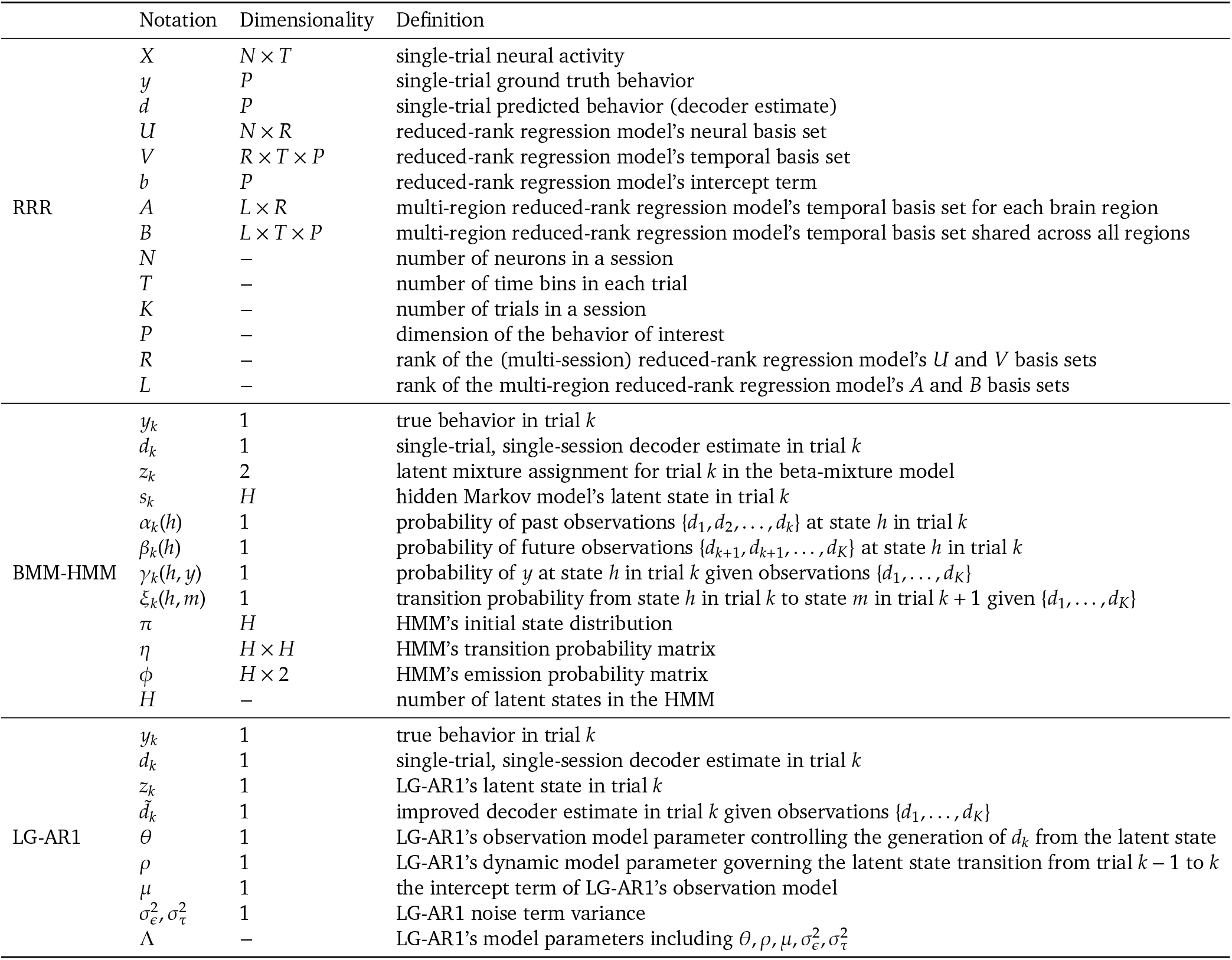
Table of notation.

|  | Notation | Dimensionality | Definition |
| --- | --- | --- | --- |
| RRR | $X$ | $N \times T$ | single-trial neural activity |
| | $y$ | $P$ | single-trial ground truth behavior |
| | $d$ | $P$ | single-trial predicted behavior (decoder estimate) |
| | $U$ | $N \times R$ | reduced-rank regression model's neural basis set |
| | $V$ | $R \times T \times P$ | reduced-rank regression model's temporal basis set |
| | $b$ | $P$ | reduced-rank regression model's intercept term |
| | $A$ | $L \times R$ | multi-region reduced-rank regression model's temporal basis set for each brain region |
| | $B$ | $L \times T \times P$ | multi-region reduced-rank regression model's temporal basis set shared across all regions |
| | $N$ | – | number of neurons in a session |
| | $T$ | – | number of time bins in each trial |
| | $K$ | – | number of trials in a session |
| | $P$ | – | dimension of the behavior of interest |
| | $R$ | – | rank of the (multi-session) reduced-rank regression model's $U$ and $V$ basis sets |
| | $L$ | – | rank of the multi-region reduced-rank regression model's $A$ and $B$ basis sets |
| BMM-HMM | $y_k$ | 1 | true behavior in trial $k$ |
| | $d_k$ | 1 | single-trial, single-session decoder estimate in trial $k$ |
| | $z_k$ | 2 | latent mixture assignment for trial $k$ in the beta-mixture model |
| | $s_k$ | $H$ | hidden Markov model's latent state in trial $k$ |
| | $\alpha_k(h)$ | 1 | probability of past observations $\{d_1, d_2, \dots, d_k\}$ at state $h$ in trial $k$ |
| | $\beta_k(h)$ | 1 | probability of future observations $\{d_{k+1}, d_{k+2}, \dots, d_K\}$ at state $h$ in trial $k$ |
| | $\gamma_k(h, y)$ | 1 | probability of $y$ at state $h$ in trial $k$ given observations $\{d_1, \dots, d_K\}$ |
| | $\xi_k(h, m)$ | 1 | transition probability from state $h$ in trial $k$ to state $m$ in trial $k + 1$ given $\{d_1, \dots, d_K\}$ |
| | $\pi$ | $H$ | HMM's initial state distribution |
| | $\eta$ | $H \times H$ | HMM's transition probability matrix |
| | $\phi$ | $H \times 2$ | HMM's emission probability matrix |
| | $H$ | – | number of latent states in the HMM |
| LG-AR1 | $y_k$ | 1 | true behavior in trial $k$ |
| | $d_k$ | 1 | single-trial, single-session decoder estimate in trial $k$ |
| | $z_k$ | 1 | LG-AR1's latent state in trial $k$ |
| | $\tilde{d}_k$ | 1 | improved decoder estimate in trial $k$ given observations $\{d_1, \dots, d_K\}$ |
| | $\theta$ | 1 | LG-AR1's observation model parameter controlling the generation of $d_k$ from the latent state |
| | $\rho$ | 1 | LG-AR1's dynamic model parameter governing the latent state transition from trial $k - 1$ to $k$ |
| | $\mu$ | 1 | the intercept term of LG-AR1's observation model |
| | $\sigma_\epsilon^2, \sigma_\tau^2$ | 1 | LG-AR1 noise term variance |
| | $\Lambda$ | – | LG-AR1's model parameters including $\theta, \rho, \mu, \sigma_\epsilon^2, \sigma_\tau^2$ |

#### Closed-form solution for reduced-rank regression model

In practice, the parameters of the reduced-rank regression model can be learned via automatic differentiation. In the special case of a linear model, a closed-form solution can be derived to improve computational efficiency and provide theoretical insight. For notational simplicity, we omit the session index *i* and represent the neural activity and behavior across all trials as *X* ∈ ℝ^*N*×*T*^ and *d* ∈ ℝ^*P*^, assuming there is only one trial in the data. To avoid dealing with the intercept term *b* from Equation 2, we use the centered neural activity and behavior matrices 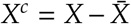 and 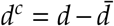, where 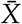 and 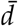 represent the trial averages.

Traditionally, a classical version of reduced-rank regression exists with a well-known closed-form solution (Izenman, 1975). This approach factorizes the full-rank regression weights into two lower-rank components:

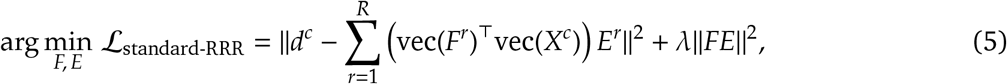

where *F* ∈ ℝ^*N*×*T*×*R*^ defines *R* basis functions (*F*^*r*^ ∈ ℝ^*N*×*T*^) over both the neural and temporal dimensions of the input neural activity *X*^*c*^, while *E* ∈ ℝ^*R*×*P*^ maps from the low-rank latent space to behavior *d*^*c*^, with *E*^*r*^ ∈ ℝ^*P*^ denoting the rank-*r* component.

Our proposed formulation (Equation 6) follows the same principle of low-rank decomposition, but introduces additional structure to explicitly separate spatial (neural) and temporal components of the regression weights. Instead of learning unconstrained spatiotemporal bases *F*^*r*^ ∈ ℝ^*N*×*T*×*R*^ and behavior-specific bases *E* ∈ ℝ^*R*×*P*^, we decompose the regression weights into a low-rank neural basis *U* ∈ ℝ^*N*×*R*^ and a low-rank temporal basis *V* ∈ ℝ^*R*×*T*×*P*^. This parameterization also allows us to share temporal structure across sessions via the temporal basis, while retaining session-specific variability through the neural basis. Specifically, our model solves the following objective:

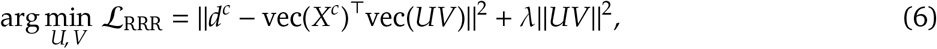

where the operator vec(·) flattens a matrix or tensor into a vector and || · ||^2^ denotes the Frobenius norm. Regularization is included via λ, though in practice it is applied as weight decay during optimization and is therefore not explicitly shown in Equations 1–4.

To obtain the solution of *V*, we differentiate ℒ_RRR_ (Equation 6) with respect to *V*:

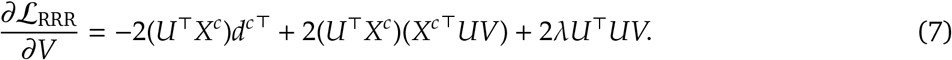

By setting the gradient in Equation 7 to 0, we obtain the closed-form solution of *V*:

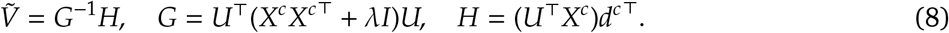

(Further details are provided below.) By substituting *V* with 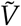 in Equation 6 (Zheng et al., 2015), the objective function simplifies to

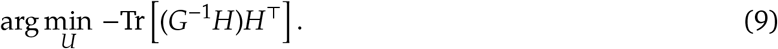

The solution of *U* is then derived by solving the following problem:

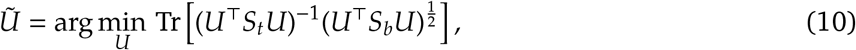

where

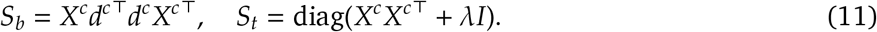

(Further details are provided below.) The resulting expression in Equation 10 constitutes a classical generalized Rayleigh quotient problem (Courant and Hilbert, 1953; Fukunaga, 1990), whose optimal solution is given by the top *R* generalized eigenvectors of the matrix pair (*S*_*b*_, *S*_*t*_). That is,

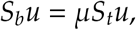

where the optimal columns of 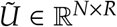 are formed from the eigenvectors *u* associated with the top *R* eigenvalues µ.

We can equivalently express Equation 10 as

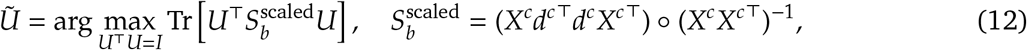

where ◦ denotes element-wise matrix multiplication. This formulation suggests that 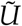 captures directions in neural space that maximize a normalized version of the neural-behavioral covariance, after correcting for neuron-specific variance. As such, 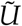provides a principled way to quantify each neuron’s contribution to decoding behavior and identifies the most influential neurons for the task. Once the optimal *U* and *V* are obtained, neural activity *X* is projected onto the learned low-rank subspace via *W* = *X*^⊤^*U*, which produces a low-dimensional representation that captures behaviorally relevant neural dynamics (Qin, 2006; Kobak et al., 2016; Sani et al., 2021). While this closed-form solution is limited to linear models, more flexible modeling frameworks using nonlinear decoders can be optimized using automatic differentiation.

#### Derivation of 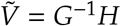 in Equation 8

To find the optimal *V*, we set the gradient in Equation 7 to 0 and obtain

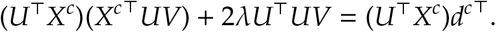

This is a linear matrix equation of the form *GV* = *H*, where

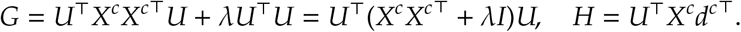

Assuming *G* is invertible, the solution is 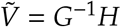 as in Equation 8.

#### Derivation of 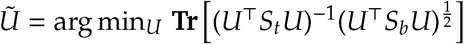 in Equation 10

For notational clarity, we omit the vectorization operator vec(·) in the remainder of this derivation. By substituting *V* with 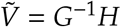 in Equation 6 (Zheng et al., 2015), we now express the objective explicitly as a function of *U*. We begin by expanding the first term in Equation 6:

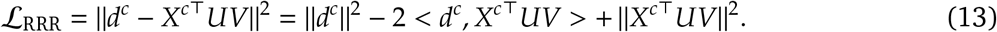

The inner product can be rewritten using the trace operator:

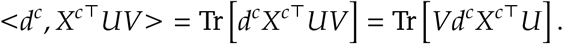

The squared norm term becomes

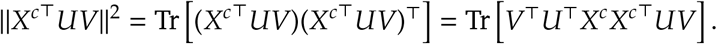

The regularization term is

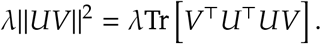

Combining these, the full loss in Equation 13 becomes

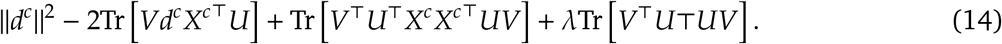

Noting that the second and third trace terms in Equation 14 can be grouped using the definition *G* = *U*^⊤^(*X*^*c*^*X*^*c*⊤^ + λ*I*)*U*, we write

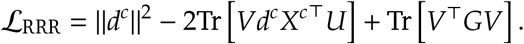

Substituting *H* = (*U*^⊤^*X*^*c*^)*d*^*c*⊤^ and *V* = *G*^−1^*H* into the expression yields:

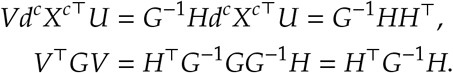

Therefore, the loss function becomes

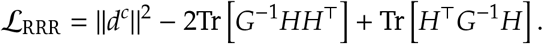

Since the two trace terms are equal (i.e., 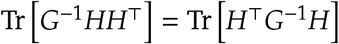), we simplify:

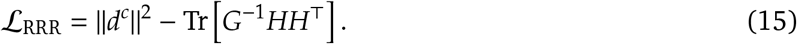

As ||*d*^*c*^||^2^ does not depend on *U*, minimizing the loss in Equation 15 is equivalent to maximizing the trace term in Equation 8:

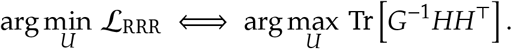

We can simplify *HH*^⊤^ and *G* further:

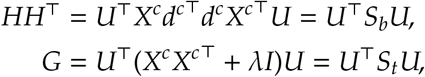

where we have defined

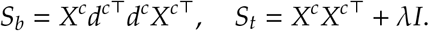

Hence, the objective becomes

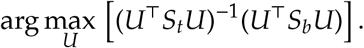

#### Connection to PCA, CCA, demixed PCA and sliced TCA

Our proposed RRR is similar to dimensionality reduction techniques like PCA and CCA, but with different objectives. As shown in Equation 12, the proposed RRR maximizes the correlation between the centered predictor *X* and the centered response *d*, as well as the variance of *d*:

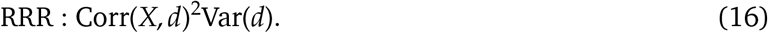

According to De la Torre (2012), PCA and CCA aim to maximize:

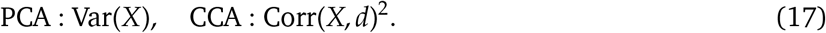

PCA captures the major variations in neural activity *X* but ignores the variations in behavior *d*, while CCA considers the correlation between *X* and *d* but doesn’t prioritize modeling the variations in *d*. The proposed RRR balances both the correlation between *X* and *d* and the variance of *d*, making it more suitable for decoding tasks where capturing behavioral variations is crucial for prediction.

The proposed RRR is also related to demixed-PCA (Kobak et al., 2016), which minimizes the loss

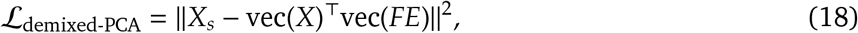

where *X* is the centered data matrix, with each row representing the neural activity of each neuron across all trials and task conditions. The reconstruction target, *X*_*s*_, is a matrix of stimulus averages, with each data point replaced by the average neural activity for the corresponding stimulus. The solutions for *F* and *E* can be analytically obtained using the standard reduced-rank regression through singular value decompositions. The main difference is the reconstruction target: the behavior *d* in our model (Equation 6) vs. the task-condition averaged neural activity *X*_*s*_ in demixed PCA. Intuitively, demixed-PCA maximizes the correlation between the neural activity *X* and the task-condition averaged neural activity *X*_*s*_, while also maximizing the variance of *X*_*s*_.

The decomposition of the full-rank weight matrix in Equation 2 resembles sliced TCA (Pellegrino et al., 2024) when the behavior dimension *P* > 1:

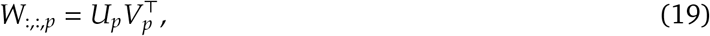

where *W* ∈ ℝ^*N*×*T*×*P*^ is the three-way regression tensor, and *U*_*p*_ ∈ ℝ^*N*×*R*^ and *V*_*p*_ ∈ ℝ^*R*×*T*^ are the low-rank factors for the *p*-th slice along the behavior dimension.

### BMM-HMM: Model details

This section presents algorithms and implementation details for various BMM-HMM model variants. The BMM-HMM model consists of a dynamic process governing transitions among discrete latent states 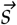 and an observation process describing the generation of decoder estimates 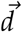 given the latent state. The dynamic model, *P*(*s*_*k*_ |*s*_*k*−1_), describes the state transition from trial *k ™* 1 to *k*, parameterized by a state transition matrix. The observation model, *p*(*d*_*k*_|*s*_*k*_) = *p*(*d*_*k*_|*z*_*k*_)*p*(*z*_*k*_|*s*_*k*_), is characterized by a beta mixture model, where *p*(*z*_*k*_ *s*_*k*_) is the emission probability at each state, *p*(*d*_*k*_|*z*_*k*_) is the observation probability, and *z*_*k*_ controls the assignment of beta distributions in the mixture.

Specifically, we assume the single-session, single-trial decoder output *d*_*k*_ = *P*(*y*_*k*_ = 1|*X*_*k*_) ∈ [0, 1] follows a mixture of beta distributions, with mixture assignment *z*_*k*_ depending on a latent state *s*_*k*_, governed by an *H*-state HMM. The data generation process for *d*_*k*_ is formulated as

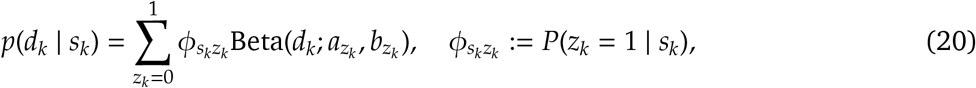

where 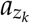 and 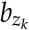 are parameters of a beta distribution. In each trial, the latent state *s*_*k*_ generates *z*_*k*_ with emission probability 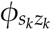, and *d*_*k*_ is drawn from a beta mixture with observation probability *p*(*d*_*k*_ *z*_*k*_), where *d*_*k*_ values cluster around 1 when *z*_*k*_ = 1 and around 0 when *z*_*k*_ = 0.

The main idea is to substitute the single-session and single-trial decoder output *d*_*k*_, which only considers information from the neural activity *X*_*k*_, with the inferred *z*_*k*_. The inferred *z*_*k*_ contains information about the underlying behavioral states deduced from the trial-to-trial correlations in 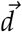. Specifically, the improved decoder output is

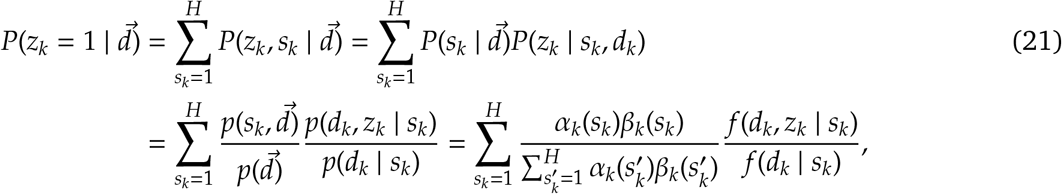

where 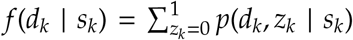, as defined in Equation 20. α_*k*_(*s*_*k*_) and β_*k*_(*s*_*k*_) are outputs from the forward and backward passes in an Expectation-Maximization (EM) algorithm, described in more depth in the next section. We focus on inferring *z*_*k*_ because it reflects the true choices made by the mice during the visual decision-making task. In contrast, *d*_*k*_ represents noisy observations influenced by these latent choices. Since our goal is to fit a latent dynamical model that recovers these underlying decisions, *z*_*k*_ is the primary target of inference.

### EM algorithm for BMM-HMM

The EM (Baum–Welch) algorithm is used for iterative HMM parameter estimation. Each iteration consists of the following Expectation and Maximization steps:

- **(E step)** Let *k* index trial, *z* ∈ {0, 1} index the beta mixture component and *h, m* ∈ {1, …, *H*} index the state. For all component and state pairs, we recursively compute the forward and backward probabilities α_*k*_(*h, z*) and β_*k*_(*h, z*), defined below. We then compute the component and state occupation probabilities γ_*k*_(*h, z*) and ξ_*k*_(*h, m*).
- **(M step)** Using the estimated γ_*k*_(*h*) and ξ_*k*_(*h*), we then update the model parameters, including the transition probabilities η_*hm*_ and the emission probabilities ϕ_*hz*_ of the HMM, and the parameters of the beta mixture *a*_*z*_, *b*_*z*_.

#### Forward pass

We define the probability of observing the sequence of decoder outputs 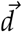 being in state *h* in trial *k* as

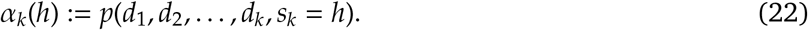

The pseudo-code for the iterative computation of α_*k*_(*h*) is:

- *Initialization* α_1_(*h*) = π_0_(*h*) *f* (*d*_1_ | *h*) ∀ 1*hH*.
- *Recursion* 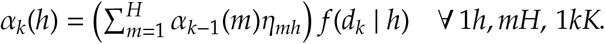
- *Termination* 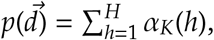

where π_0_ is a vector containing the initial probabilities for each of the *H* hidden states.

#### Backward pass

The probability of future observations given that the HMM is in state *h* in trial *k* is

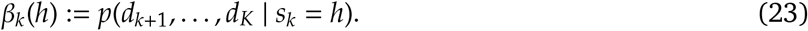

The pseudo-code for the iterative computation of β_*k*_(*h*) is:

- *Initialization* β_*K*_(*h*) = 1 ∀ 1*hH*.
- *Recursion* 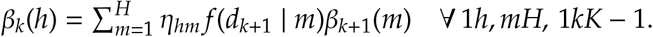
- *Termination* 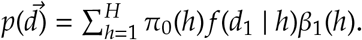

#### Forward-backward

The state occupation probability γ_*k*_(*h*) is

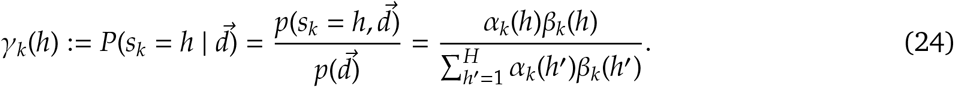

The component and state occupation probability γ_*k*_(*h, z*) is the probability of component *z* at state *h* in trial *k* given the whole observation sequence 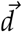:

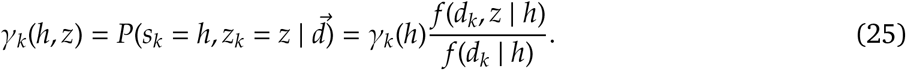

We then estimate ξ_*k*_(*h, m*), the probability of transitioning from state *h* to *m* given all observations 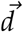:

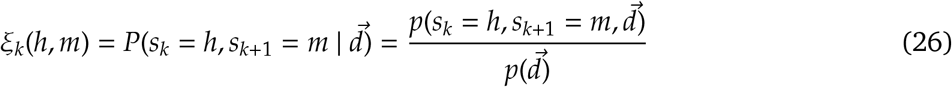

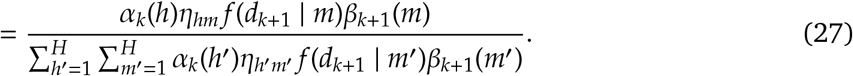

For the M step, we update the transition and emission probabilities according to

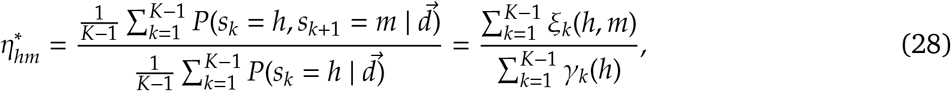

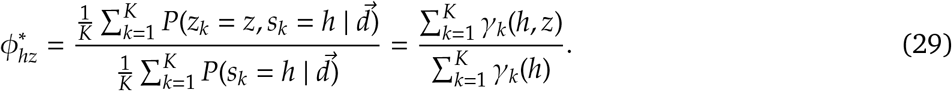

We then update the parameters of the BMM (*a*_0_, *a*_1_, *b*_0_, *b*_1_) by maximizing the expected log-likelihood. First, we write down the likelihood of the BMM as

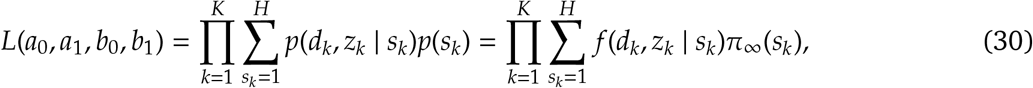

where π_∞_ represents the equilibrium probability for each of *H* hidden states, which can be computed using the estimated transition probabilities. The conditional distribution is subsequently determined by

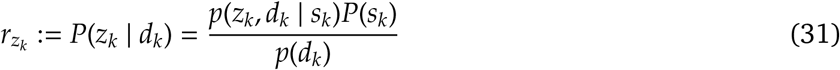

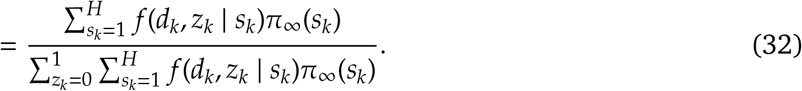

Finally, the expected log-likelihood of the BMM is

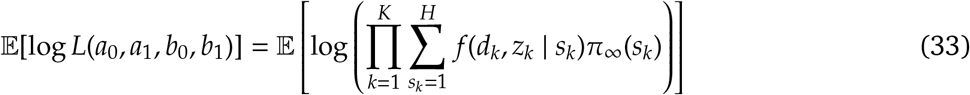

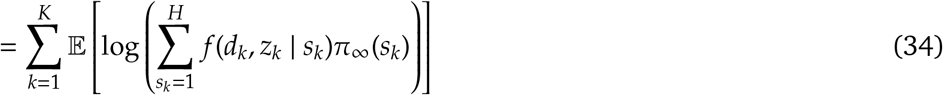

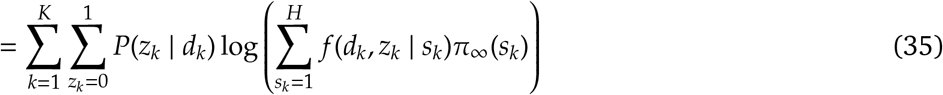

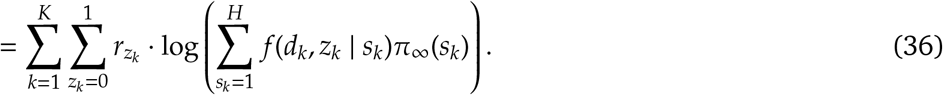

In practice, we find 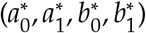 that maximize the quantity in Equation 36 through numerical optimization.

### Oracle BMM-HMM

In each session, the oracle BMM-HMM substitutes the ground truth observed behaviors 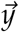 for 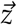, treating 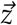 as a known quantity. This allows us to learn the underlying data-generating mechanism that produces the decoder outputs 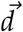. The process consists of the following steps:

1. Train a discrete-state HMM on the ground truth observed behaviors 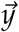 to estimate the oracle model parameters, including transition probabilities η_*hm*_ and emission probabilities ϕ_*hz*_ for each session.
2. Apply a BMM to the decoder outputs 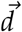, treating the mixture assignment variable 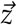 as a known quantity by substituting 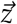 with the ground truth observed behaviors 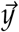. This step provides the correct assignment of mixture components. The learned oracle BMM parameters, (*a*_0_, *a*_1_, *b*_0_, *b*_1_), capture the true probabilistic relationship between 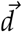 and 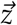.
3. Use the learned oracle model parameters to initialize and fit the BMM-HMM using the EM algorithm described in the section “EM algorithm for BMM-HMM” for the corresponding session. During model fitting, fix the oracle parameters (η_*hm*_, ϕ_*hz*_, *a*_0_, *a*_1_, *b*_0_, *b*_1_). Although the model parameters remain fixed, we infer the latent neural states *s*_*k*_ and the latent choices *z*_*k*_ based on the observed baseline decoder output *d*_*k*_.

This procedure allows us to deduce the latent behavioral states 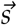 and latent behaviors 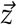 as if we know the true data generation process.

### Learning empirical priors of state-space model parameters

To learn empirical priors for the multi-session BMM-HMM, we fit a variational HMM (Gruhl and Sick, 2016) to the ground truth observed behavior 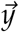 from non-target sessions. This allows us to learn an empirical prior of the trial-to-trial correlations inherent in the true behavioral data. We impose Dirichlet priors on the initial state distribution π_0_, rows of the transition probability matrix η_*h*·_, and rows of the emission probability matrix ϕ_*h*·_ as follows:

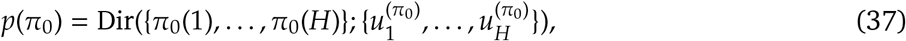

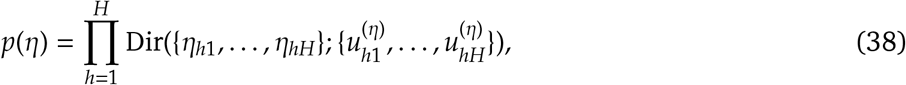

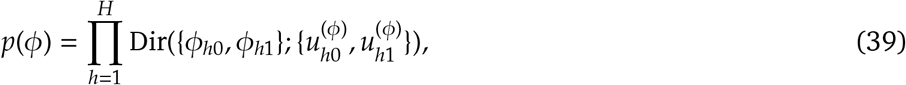

where 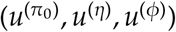 are the Dirichlet distribution concentration parameters, learned by fitting a variational HMM on the ground truth observed behaviors 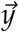 from the training sessions using the Python package *hmmlearn*. The resulting posterior distributions serve as priors for the multi-session BMM-HMM parameters, constraining their updates during the EM algorithm’s M step.

To set empirical priors for the BMM parameters, we assume *d*_*k*_ follows a mixture of beta distributions from the exponential family, expressed as:

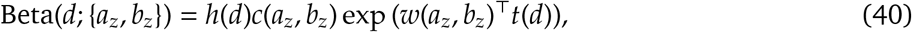

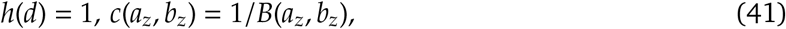

where

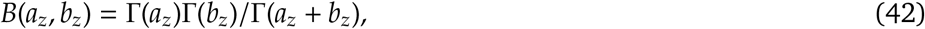

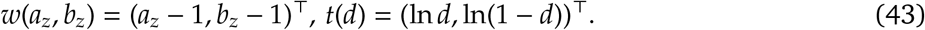

For exponential family members, the conjugate prior is

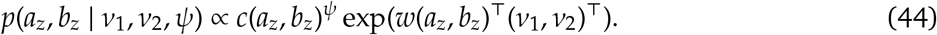

**Figure S1:**
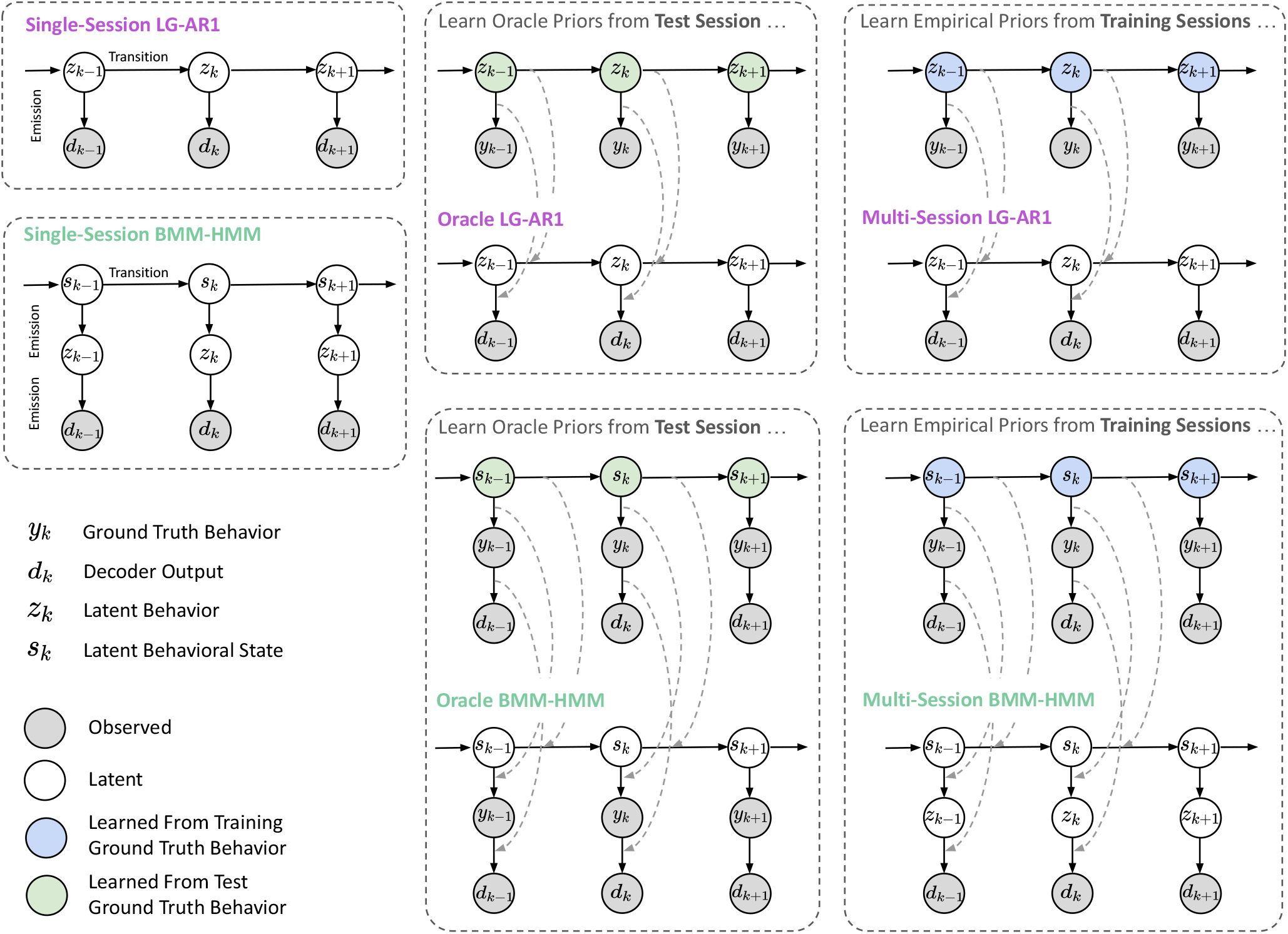
Graphical models for single-session, oracle, and multi-session LG-AR1 and BMM-HMM models. The single-session LG-AR1 and BMM-HMM models learn parameters directly from the test session. In contrast, the oracle versions of these models use a two-step learning process. They first derive oracle priors from the test set’s ground truth behavior, then use these priors to guide parameter learning on the test data, as indicated by gray arrows. The multi-session variants follow a similar approach. They learn empirical priors from the training data’s ground truth behaviors, which then constrain parameter learning on the test set, also shown by gray arrows. This approach allows the multi-session models to leverage information from multiple training sessions while adapting to the current test data.

Therefore, a suitable conjugate prior distribution for (*a*_*z*_, *b*_*z*_) is

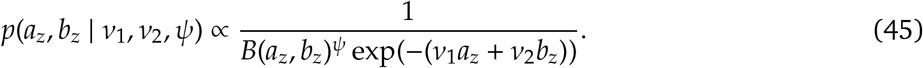

Setting the natural conjugate prior ψ parameter to zero yields independent exponential priors for (*a*_*z*_, *b*_*z*_), which have proven effective empirically. We apply a hierarchical BMM on the decoder outputs 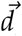, using the Python package *pymc3*. We assume that the mixture assignment 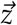 can be empirically determined a priori, and substitute 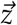 with the observed behaviors 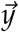 from the training sessions. The posterior distributions for 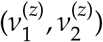 then serve as priors for the multi-session BMM-HMM parameters, constraining their updates during the EM algorithm’s M step.

### Multi-session BMM-HMM

Following the approach in Equation 37-39, we impose Dirichlet priors on the BMM-HMM dynamic parameters (π_0_, η_*h*·_, ϕ_*h*·_). We modify the EM algorithm in the section “EM algorithm for BMM-HMM” by using Maximum A Posteriori (MAP) estimation (Bassett and Deride, 2019) to learn the posterior distributions of these parameters. The E step remains unchanged, while the M step incorporates the new prior terms when updating the HMM parameters with fixed latent *s*_*k*_ and *z*_*k*_. The posterior means of the HMM parameters become

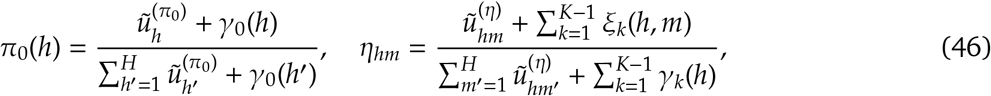

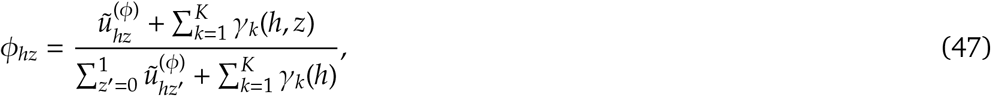

where 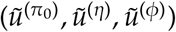 are the posterior concentration parameters from fitting the variational HMM on the training sessions. When updating BMM parameters, we add the Dirichlet prior term log *p*(π_0_, η, ϕ) to the complete-data log-likelihood in Equation 33 and solve for (*a*_0_, *a*_1_, *b*_0_, *b*_1_) that maximize this new objective function.

We constrain BMM parameters (*a*_0_, *a*_1_, *b*_0_, *b*_1_), using empirical priors, 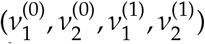, learned from the training sessions; see details in the section “Learning empirical priors of state-space model parameters”. Incorporating the log-prior term (Equation 45) into the complete log-likelihood involves adding the following penalty to the right-hand side of Equation 36:

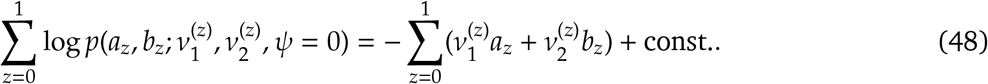

Numerically solving the penalized objective yields MAP estimates for the BMM parameters instead of the standard maximum likelihood estimation (MLE) solutions.

### LG-AR1: Model details

For scalar-valued *y*_*k*_ ∈ ℝ, we assume the decoder output *d*_*k*_ ∈ ℝ linearly depends on the latent behavior *z*_*k*_ ∈ ℝ. To incorporate trial-to-trial correlations, the transitions of *z*_*k*_ between trials are modeled using a first-order autoregressive process. The objective aligns with that of a Kalman smoother (Welch et al., 1995), which is to infer the state of a dynamical system (*z*_*k*_) given a sequence of noisy observations (*d*_*k*_). The formal data generating model is described as

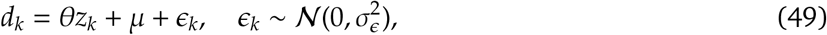

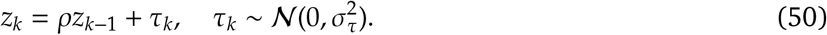

Intuitively, as the *first-order autocorrelation coefficient* ρ approaches 1, *z*_*k*_ in the current trial is expected to exhibit minimal deviation from *z*_*k*−1_ in the preceding trial, as per Equation 50. As the *linear coefficient of the Gaussian observation model* θ approaches 1, *d*_*k*_ is expected to closely track the pattern of *z*_*k*_ according to Equation 49. µ is the *bias of the Gaussian observation model*. In practice, the values of θ and ρ are determined by fitting the LG-AR1 model to the observed 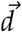.

Similar to BMM-HMM, the main idea is to replace the original decoder estimate *d*_*k*_, based solely on neural activity *X*_*k*_, with a smoothed estimate 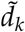 derived from the inferred latent state 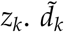 incorporates trial-to-trial correlations from 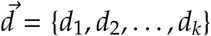, as 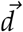 is used to infer the latent states 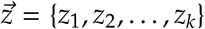. This process potentially improves 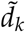’s accuracy over the original *d*_*k*_ in estimating the true behavior. While 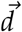 is used for model fitting and latent state inference, 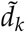 is the improved (smoothed) decoder estimate for the held-out trial *k* given the entire 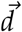. To obtain 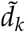, we sample from its posterior predictive distribution

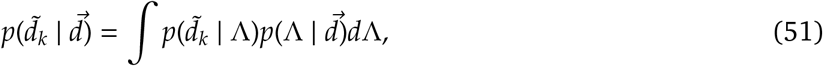

after placing prior distributions on the model parameters 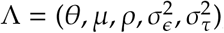, which can be estimated using Markov chain Monte Carlo (MCMC) sampling (Brooks, 1998).

In Equation 51, we use 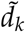, the posterior mean of *d*_*k*_, because empirically it leads to better recovery of the prior variable compared to using the inferred *z*_*k*_. We believe this is due to the fact that the prior we aim to decode is itself the output of another model (Findling et al., 2023), which estimates the “true” prior from data and thus contains noise. Ideally, we would compare the latent *z*_*k*_ directly to the true, unobserved prior, but since that is inaccessible, we rely on an estimated version from a statistical model (Findling et al., 2023). In this context, 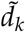 serves as a proxy.

To fit LG-AR1 on single-session data, we use a Bayesian approach, treating model parameters 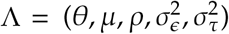 as random variables with joint prior *p*(Λ):

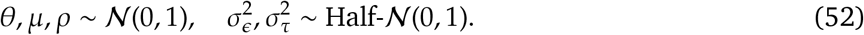

In practice, we use the Python package *pymc3* to fit the hierarchical LG-AR1 and learn the posterior distribution of session-specific parameters Λ via MCMC sampling.

To implement the multi-session LG-AR1, we begin by learning the dynamic model parameters 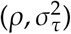. This estimation is performed using the observed behaviors 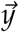 from the training sessions, under the assumption that these dynamic model parameters can be empirically determined a priori. Next, we estimate observation model parameters 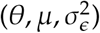 using decoder outputs 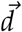 and corresponding observed 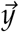 from training sessions. After estimating model parameters from the training data, we use the posterior means of these multi-session LG-AR1 parameters 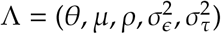 to initialize the hierarchical LG-AR1 model (Equation 49-50) for the held-out session, with Λ fixed during model fitting. For this held-out session, where true behaviors are unknown, we infer the latent behaviors *z*_*k*_ and obtain improved decoder outputs 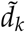 via MCMC sampling.

We also implement an oracle LG-AR1 model to emulate the ground-truth data-generating process for 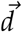. This oracle model is constructed by estimating model parameters using the ground truth observed 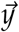 from the target session, under the assumption that the true values of the variable 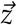 are known. For the oracle model, we learn dynamic AR1 parameters 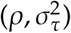 and observation model parameters 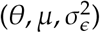 using true 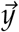 and observed 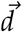 from the test session. We initialize the hierarchical LG-AR1 model using these oracle solutions and hold them fixed while inferring the latent *z*_*k*_ and improved decoder outputs 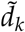, as if we know the true data-generating mechanism.

### Quantification and statistical analysis

#### Data processing

##### IBL trial-aligned dataset

For choice, we align trials to the stimulus onset, considering neural activity from 0.5 seconds before to 1.5 seconds post-onset. For prior, we also align trials to the stimulus onset, including neural activity from 0.6 seconds to 0.1 seconds pre-onset. The prior represents the mice’s estimate of the stimulus side probability. We use the same decoding window as in the previous study (Findling et al., 2023), focusing on the period with minimal wheel movements. Within each trial, we segment neural activity into 50-millisecond non-overlapping time bins. For each time bin, we bin spike counts using all neurons, sorted by Kilosort 2.5 (Pachitariu et al., 2016), from each session. For dynamic behaviors (wheel speed and whisker motion energy), we select the first movement onset as the alignment event, and decode the behavior starting at the alignment event and ending at 1 second after the alignment event. The neural activity within each trial is binned into non-overlapping 20 ms bins. For each time bin, we similarly bin spike counts using all neurons from each session.

##### IBL trial-unaligned dataset

We focus only on wheel speed and whisker motion energy, as variables like choice and prior are only defined during experimental trials, which are periods when stimuli are presented and mice respond for rewards. Unlike the trial-aligned dataset, we do not align neural activity and behavior to a specific event onset. Instead, we segment the continuous Neuropixels recordings into 2-second snippets to construct the trial-unaligned dataset for each session. These snippets cover intervals both during and outside experimental trials, capturing the repetitive, structured behaviors within trials as well as the more spontaneous behaviors outside of trials.

##### Allen dataset

To use reduced-rank regression models, we decode 3 behavior variables: Gabors, static gratings, and running speed. The Gabors variable has 3 discrete orientation classes, while the static gratings variable has 6. Running speed, in contrast, is a continuous trial-unaligned variable recorded during periods of spontaneous behavior. For Gabors and static gratings, we align trials to the respective stimulus onset and extract neural activity within a 200 ms window around the onset. For running speed, we divide the continuous recording into 1-second snippets. For all variables, we use a time bin size of 10 ms to bin neural activity.

To apply BMM-HMM, we also decode the licks. In the Allen datasets, mice view a continuous stream of briefly presented visual stimuli and receive water rewards for correctly detecting changes in stimulus identity. To relate neural activity to behavior, the timing of behavioral responses, licks, is also recorded. Each Neuropixels recording session is divided into five stimulus blocks, but licking occurs exclusively during the “active” period, which lasts approximately 2000 to 3000 seconds. During this period, each visual stimulus is updated roughly every 0.2 seconds. We segment the active period into fixed 0.2-second intervals and labeled each interval as 1 if at least one lick occurred, and 0 otherwise. For each interval, we use a time bin size of 10 ms. For each session, we decode licks using neural activity from the recorded subset of regions of interest, DG, CA1, CA3, VISp, VISa, LP, and TH (the medial division of the posterior parahippocampal cortex). For example, if a session records from DG, CA1, and LP, all three are used in the decoder. Regions outside this set are excluded from the analysis, even if recorded. Region naming follows the IBL brain atlas conventions.

##### Primate random target dataset

This dataset consists of 15 minutes of continuous reaching behavior, manually segmented into 1351 600 ms “trials.” It includes recordings from 130 neurons along with simultaneously tracked hand kinematics. We use the preprocessed version of the dataset provided by Pei et al. (2021), and apply a 20 ms bin size to discretize neural activity and behavior into 30 time bins per trial. Our decoding target is finger velocity, represented by velocity components along both the X and Y axes.

### Hyperparameter selection

For static variables (choice, prior, Gabors, static gratings), the target behavior in each trial is decoded using the full neural activity from that trial. For dynamic behaviors (wheel speed, whisker motion energy, running speed, finger velocity), the target behavior at each time bin *t* is decoded using the entire neural activity across all time bins within the corresponding trial. For multi-session models applied to the IBL dataset, we use both multi-session and multi-region reduced-rank regression models to decode static and dynamic behaviors. In contrast, for the Allen and primate random target datasets, we do not include multi-region reduced-rank regression models, as our objective is not to perform region-specific decoding for these datasets. For discrete behavior variables (e.g., choice and licks), we employ both single- and multi-session BMM-HMM models. To decode prior, we use both single- and multi-session LG-AR1 models. Decoder performance is evaluated using accuracy and AUC for discrete variables, Pearson correlation for prior, and *R*^2^ for dynamic behaviors.

For our linear baselines, we use L2-regularized logistic regression for discrete-valued variables and ridge regression for prior and dynamic behavioral variables. Regularization strength is selected via *scikit-learn*’s GridSearchCV function, from the set {10^−4^, 10^−3^, 10^−2^, 10^−1^, 10^0^, 10^1^, 10^2^, 10^3^, 10^4^}. Both reduced-rank regression models and MLPs are trained using gradient descent in PyTorch, with the AdamW optimizer and a cosine annealing learning rate scheduler. All models are trained for 2000 epochs with a batch size of 16, and the best model, determined by the lowest validation loss, is selected for test set decoding. For single-session MLP and reduced-rank regression models, hyperparameter search is performed over the value ranges in Table 2, using *Ray Tune* in Python. We randomly sample 30 models from the specified range for this search. For multi-session and multi-region reduced-rank regression models, extensive hyperparameter tuning is not required as for the single-session models, since the model is less prone to overfitting. As a result, we only perform a search over the rank of the reduced-rank regression models, while fixing the learning rate at 0.01 and weight decay at 1, based on results from pilot studies.

**Table 2:**
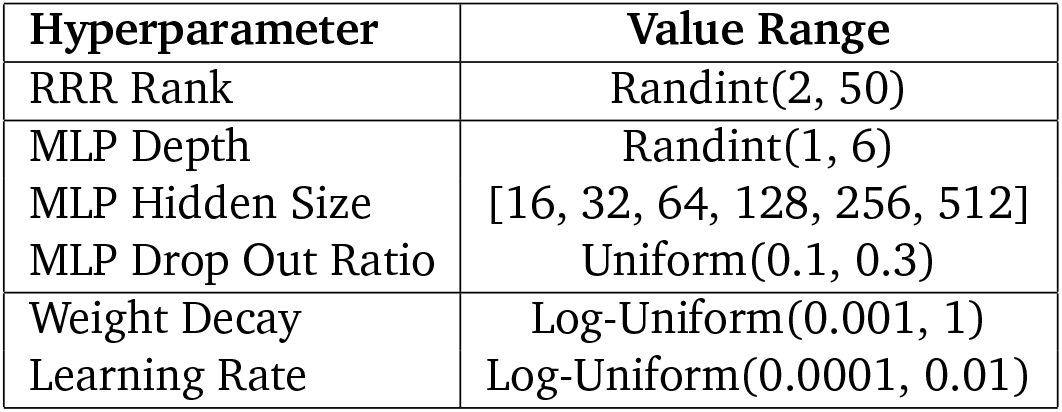
The range of possible model and optimizer hyperparameters from which *Ray Tune* randomly samples combinations.

**Table 3:**
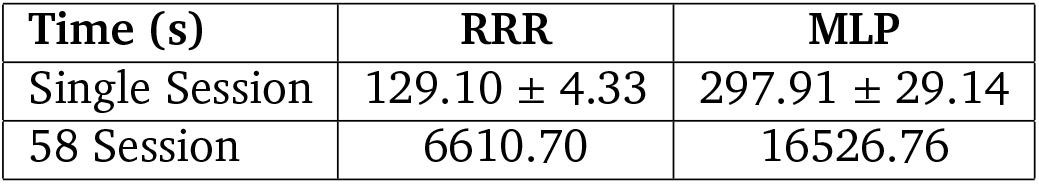
Computation time (in seconds) for single-session and multi-session reduced-rank regression models (RRR) and MLPs. For single-session models, we report the mean and standard error of training times across 58 individual sessions. For multi-session models, we report the total time required to train a single model on all 58 sessions.

| Time (s) | RRR | MLP |
| --- | --- | --- |
| Single Session | 129.10 ± 4.33 | 297.91 ± 29.14 |
| 58 Session | 6610.70 | 16526.76 |

**Figure S2:**
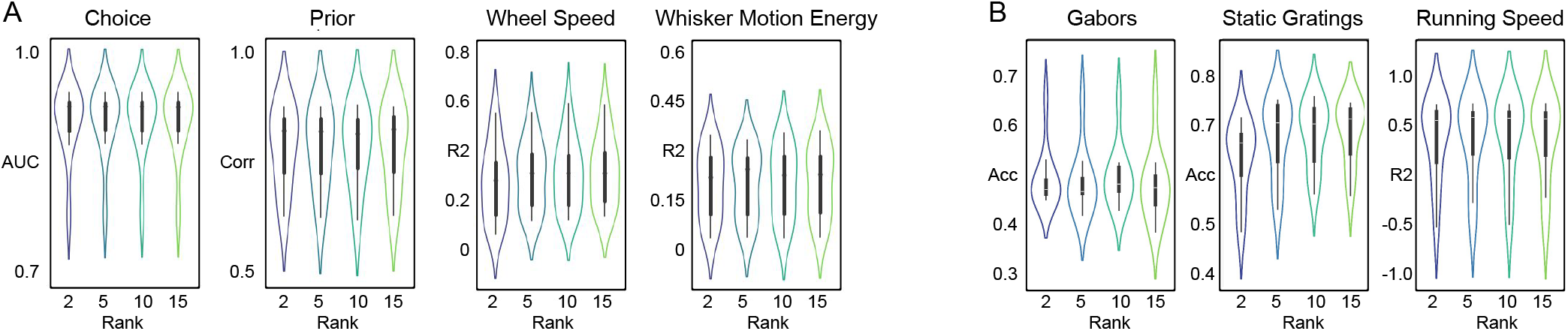
Effects of model rank on decoding performance across different behavior variables. For each behavior variable in the IBL (**A**) and Allen (**B**) dataset, the violin plots display performance metric distributions across 10 sessions when decoding with different model ranks. Although all behaviors are adequately predicted using lower model ranks, wheel speed and static gratings exhibit improved decoding performance at higher ranks.

### Recording hardwares, task structures and implementation guidelines

#### Reduced-rank regression models

These models are compatible with any electrophysiological recording hardwares that output single-spike data, including both high-density Neuropixels probes and lower-density Utah arrays. Regardless of the number of recorded neurons, reduced-rank regression models consistently demonstrate strong decoding performance and serve as a reliable alternative to traditional linear models across datasets. For example, the IBL dataset contains an average of 676 neurons per session, ranging from approximately 300 to over 2000 neurons. In comparison, the Allen dataset has an average of 690 neurons per session, with a minimum of 415 and a maximum of 1005 neurons. Even in the primate random target dataset, which contains only 130 neurons, the reduced-rank regression model outperforms the linear baseline (Figure 8 J-L). Importantly, these models are not constrained by task structure: they can be applied to both highly structured, trial-based paradigms as well as spontaneous, freely behaving tasks (Figure 8 A-C). reduced-rank regression models are capable of decoding both discrete and continuous static variables (e.g., choice and prior), as well as dynamic variables (e.g., wheel speed). Moreover, their performance is generally robust to the model rank, which is the main hyperparameter of the model. We find that the effect of rank on decoding performance is consistent across the IBL and Allen datasets (Figure S2).

#### BMM-HMM and LG-AR1 Models

These models can be used with the same electrophysiological recording hardwares as the reduced-rank regression models. The BMM-HMM is designed for decoding discrete-valued behavior variables, such as choice. While its default formulation is best suited for binary variables, it can be extended to multi-class variables using one-vs-all classification. In addition, users need to consider the nature of the behavior task, i.e., whether there is “block”-like behavioral structure in the data to begin with. For example, previous studies have shown that choice probabilities can vary across trials within a session, either due to block-based stimulus sequences in the IBL task design (IBL et al., 2022), or due to shifts in internal states, as shown in Ashwood et al. (2022). Because BMM-HMM infers latent internal states underlying observed behavioral outcomes, such as when a mouse perceives a stimulus and responds with a choice, it is not applicable to visual stimuli like Gabors and static gratings, which are not behavioral responses. The LG-AR1 model is suited for scalar-valued variables like prior. However, both models are flexible: users can modify the latent dynamical and observation models to accommodate different types of behavior variables. For example, replacing the beta mixture model in the BMM-HMM with a Gaussian mixture model allows it to model scalar-valued variables. Similarly, the LG-AR1 model can be extended to vector-valued variables, such as wheel speed or whisker motion energy, by substituting the univariate Gaussian observation model with a multivariate Gaussian and using a vector autoregressive latent dynamical model.

### Computation time comparison

We evaluate the computational efficiency of the proposed reduced-rank regression model in comparison to baseline MLP models. Specifically, we measure the real-time training duration for both single-session and multi-session RRR and MLP using the Allen visual coding dataset. For single-session models, each model is trained on one of the 58 sessions, and the average computation time is calculated across all sessions. For multi-session models, a single model is trained jointly on all 58 sessions, and the total training time is recorded. To ensure a fair comparison, all models are trained for 2000 epochs on the same hardware: a single NVIDIA RTX 3060 GPU.

We observe that training the MLP model takes approximately 2.5 times longer than training the RRR model, in both single-session and multi-session settings. The single-session reduced-rank regression model is highly efficient, requiring only about 2 minutes to complete training for 2000 epochs on a single session using a single GPU. The multi-session RRR model is also highly efficient, requiring less than 2 hours to train for 2000 epochs on the full 58-session dataset. While a direct comparison in terms of data size and experimental setup is not explored in this work, training a 100-session POYO model (Azabou et al., 2023) using 8 NVIDIA A40 GPUs reportedly takes approximately 2 days for 400 epochs. This underscores the computational efficiency of the multi-session RRR model relative to more resource-intensive transformer-based approaches.

### Assessing statistical significance of decoding improvements via bootstrap resampling

In Section 4.1 “Learning behaviorally relevant neural variations across sessions”, we utilize a bootstrap resampling approach to assess the statistical significance of the decoding improvements achieved by the multi-session RRR compared to the single-session RRR. Specifically, we calculate the difference in Adjusted Rand Index (ARI) and decoding accuracy between the two models across the same set of 10 sessions. To evaluate whether this observed difference is statistically meaningful, we generate a null distribution by resampling the computed ARI and accuracy values with replacement. For each of 10,000 bootstrap samples, we randomly sample data points with replacement from the 10 sessions and compute the ARI and accuracy difference between the models on each resampled dataset. This results in empirical null distributions under the assumption that any metric difference arises from sampling variability. We then estimate a nonparametric *p*-value by calculating the proportion of bootstrap samples in which the absolute performance difference exceeds the observed one.

### Decoding frequency components of behavior

Observing that whisker motion energy exhibits higher-frequency fluctuations than the smoother wheel speed (Figure S3A and E), we hypothesize that decoders may still effectively capture slower behavioral components even if they fail to fully reconstruct the entire signal. To answer this question, we decompose the behavioral signals into orthogonal components using both Fourier analysis and PCA, and evaluate how well different baseline decoders reconstruct each component.

To determine the amount of behavioral variance explained at each frequency, we compute the power spectral density (PSD) of the observed behavior, predicted behavior, and prediction error, following the approach from Warland et al. (1997). As shown in Figure S3B and F, the PSDs for both observed and predicted behaviors, as well as the corresponding prediction error, drop off steeply at higher frequencies. Beyond 5 Hz, the decoder captures little to no behavioral information, and most of the variation in the behavioral signals lies in the lower frequency range. We also apply PCA to the observed behavior and project the observed, predicted, and error signals onto the resulting principal components (PCs). As shown in Figure S3 B and F, the first 10 PCs account for most of the behavioral variance, and the reduced-rank regression model primarily captures information contained in these dominant components.

**Figure S3:**
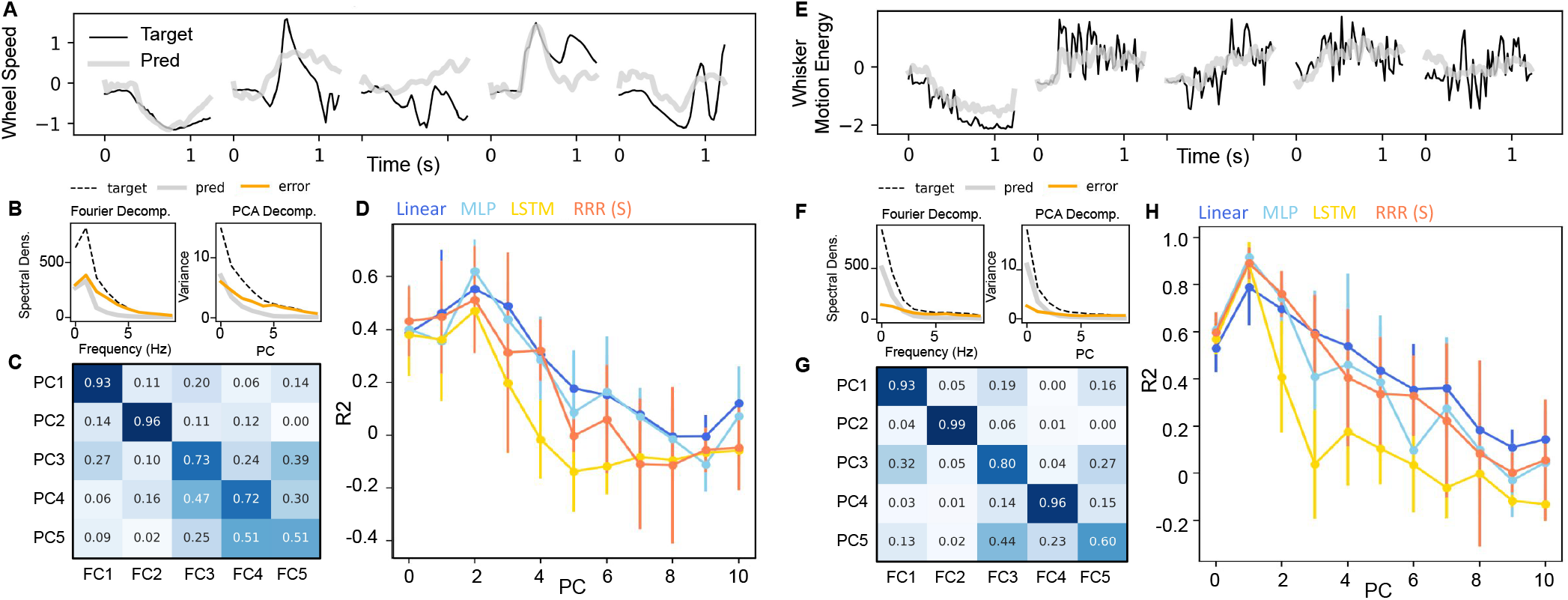
Evaluating the reduced-rank regression model against baseline decoders in capturing the primary components of the target behaviors. **(A, E)** Examples of actual (“Target”) and predicted (“Pred”) behaviors for wheel speed and whisker motion energy from the reduced-rank regression model across five selected trials. Whisker motion energy exhibits higher-frequency fluctuations compared to wheel speed. **(B, F)** Power spectral density (PSD) as a function of frequency is shown for the observed behavior (“target”), predicted behavior (“pred”) from the reduced-rank regression model, and decoding error (“error,” defined as target minus pred), averaged over 10 sessions. In addition, the variance explained by each principal component (PC) is plotted for the target, pred, and error. Both the low-frequency Fourier components and leading PCs account for most of the variance in the behavioral signals. **(C, G)** Heatmap showing the Pearson correlation between the first 5 PCs and the first 5 Fourier components (FCs) of the behavioral signals. A clear one-to-one correspondence is observed for the first two PCs and FCs, indicating that the leading PCs primarily capture low-frequency components in the behavior. **(D, H)** Model performance (*R*^2^) for decoding behaviors reconstructed from each principal component of the observed behavior using baseline decoders. Decoding *R*^2^ is generally higher for the leading PCs, which are associated with low-frequency Fourier components. The mean and standard deviation of *R*^2^ across 10 sessions are shown for the first 10 PCs.

A one-to-one correspondence between the first 5 principal components and the first 5 Fourier components (Figure S3 C and G) suggests that the leading PCs capture low-frequency components of the behavior. To determine whether the decoders can capture both gradual and rapid fluctuations in behavior, we extract the first 10 PCs from the observed behavior and reconstruct the behavior using each PC. We then train each baseline decoder to predict these PC-specific reconstructions from neural activity. Figure S3 D and H display the decoding *R*^2^ for each PC across all baseline models. Most decoders perform similarly when decoding low-frequency components, with the reduced-rank regression model slightly outperforming others for wheel speed, which is inherently smoother. Overall, higher decoding performance is achieved primarily for lower-frequency components.

### Assessing statistical significance of decoding improvements via null distribution

In Section 4.5 “Mapping behaviorally-relevant timescales across the brain,” we measure the increased information decoded from each region using the multi-region reduced-rank regression model compared to the baseline linear decoder. To control for potential spurious correlations, we conduct an additional experiment, following the approach in IBL et al. (2023). We generate null distributions to test the significance of our decoding results according to the procedure described in the caption of Figure S4. To determine the significance of our decoding results, we analyze brain regions PO, LP, DG, CA1, and VISa as representative examples. Figure S4 displays the adjusted scores, with the original scores for choice and prior decoding corresponding to those in Figure 7D. While the percentage increase in decoding metrics varies slightly between original and adjusted scores, the relative ranking of brain regions, based on decoding improvement, remains largely consistent. For instance, DG shows the highest improvement in decodable information for choice, both before and after null distribution adjustment. This analysis demonstrates the reliability of the decoding improvement offered by our proposed model.

**Figure S4:**
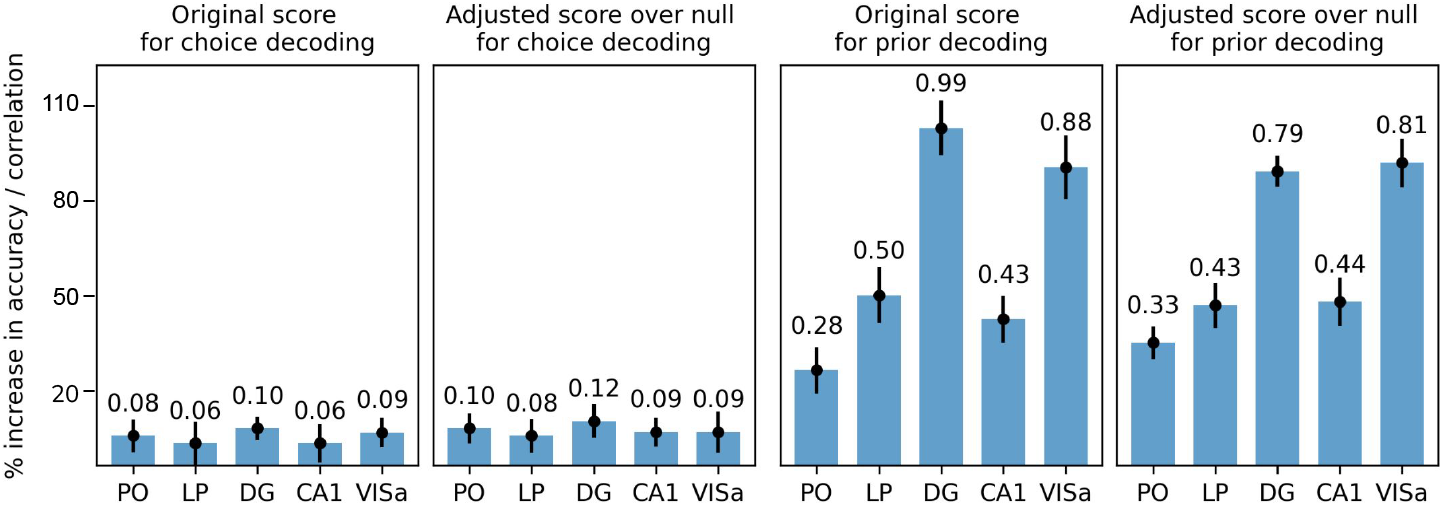
Assessing the decoding improvement achieved by multi-region reduced-rank regression model relative to null distributions generated from imposter sessions. For each session with probe insertions in PO, LP, DG, CA1, and VISa, we create 10 “imposter sessions” from behaviors (choice and prior) of other mice in different sessions, as in IBL et al. (2023). These are generated by concatenating trials across all analyzed sessions, excluding the session under consideration, then randomly selecting a chunk of *N* consecutive trials (where *N* matches the original session length) from the concatenated sessions. We obtain the original score from the real session, while the adjusted score is calculated by subtracting the decoding accuracy (or correlation) of the imposter sessions from the original score. Each bar shows the mean score from 10 imposter sessions, with error bars indicating one standard deviation of these scores.

### Relationship between LG-AR1 and post-hoc smoother, and across-session variability in BMM-HMM/LG-AR1 outputs

In Figures 4 D and H, we observe that the LG-AR1 models output a smoother version of the baseline model’s predictions. Although the LG-AR1 model is based on a latent variable framework, it can also be interpreted as a post-hoc smoother. To investigate whether applying a post-hoc smoother to denoise the baseline model’s predicted prior outperforms the LG-AR1 model, we use the Savitzky-Golay filter from the Python *scikit-learn* package. This filter smooths the decoder output by fitting successive windows of the signal with low-degree polynomials using least-squares regression. We use a window length of 17 samples and a third-order polynomial. The smoothed signal preserves key features such as peaks and slopes while reducing high-frequency noise (Figure S5B). Together, Figures S5 A and B demonstrate that the post-hoc smoother performs similarly to the single-session LG-AR1 model.

Figure 4 illustrates how BMM-HMM and LG-AR1 models enhance decoder outputs, using examples of estimated model parameters and predictions from a single session. To examine their effects across multiple sessions, we train single-session, oracle, and multi-session BMM-HMM/LG-AR1 models over 10 sessions. In Figures S5 C and D, we compare parameter estimation accuracy between the single- and multi-session models by evaluating their parameters against those of the oracle models. Using root mean squared error (RMSE) as the metric, we find that the multi-session BMM-HMM/LG-AR1 models consistently outperform the single-session models in terms of lower estimation errors. In addition to the decoded choice probabilities and priors shown in Figures 4 G and H, we present additional examples from four other sessions to examine variability in model predictions across sessions (Figure S5 E and F). We observe that the “block” structure is more clear in the decoded choice probabilities for some sessions compared to others. However, even in sessions with less apparent “block” structure, the multi-session BMM-HMM models can still exploit across-session structure to generate predictions closer to oracle BMM-HMM than single-session BMM-HMM.

### Scaling curves for multi-session reduced-rank regression models

The empirical and theoretical success of scaling laws in multi-session neuro-foundation models (Azabou et al., 2023; Ye et al., 2023; Zhang et al., 2024b; Azabou et al., 2024; Zhang et al., 2025) suggests that model performance improves predictably as the amount of data increases. These findings raise an important question: Do similar scaling laws apply to non-transformer-based, linear models, such as multi-session reduced-rank regression models? In this section, we present the results of our scaling experiments using the IBL dataset. For four behavior variables, choice, prior, wheel speed, and whisker motion energy, we train both single-session and multi-session reduced-rank regression models on data from 5, 10, 20, 40, and 73 sessions, and evaluate their performance on a fixed set of 5 test sessions. As shown in Figure S7, only whisker motion energy exhibits consistent improvement in decoding performance with additional training sessions. For choice, prior, and wheel speed, decoding performance improves up to approximately 20 ∼ 40 training sessions, after which it begins to decline. This pattern suggests that while increased training data can initially help prevent overfitting and uncover shared structure across sessions, the benefits plateau beyond a certain threshold. Overall, our findings indicate that scaling behavior in multi-session reduced-rank regression models is task-dependent, and their performance does not universally improve with more data.

**Figure S5:**
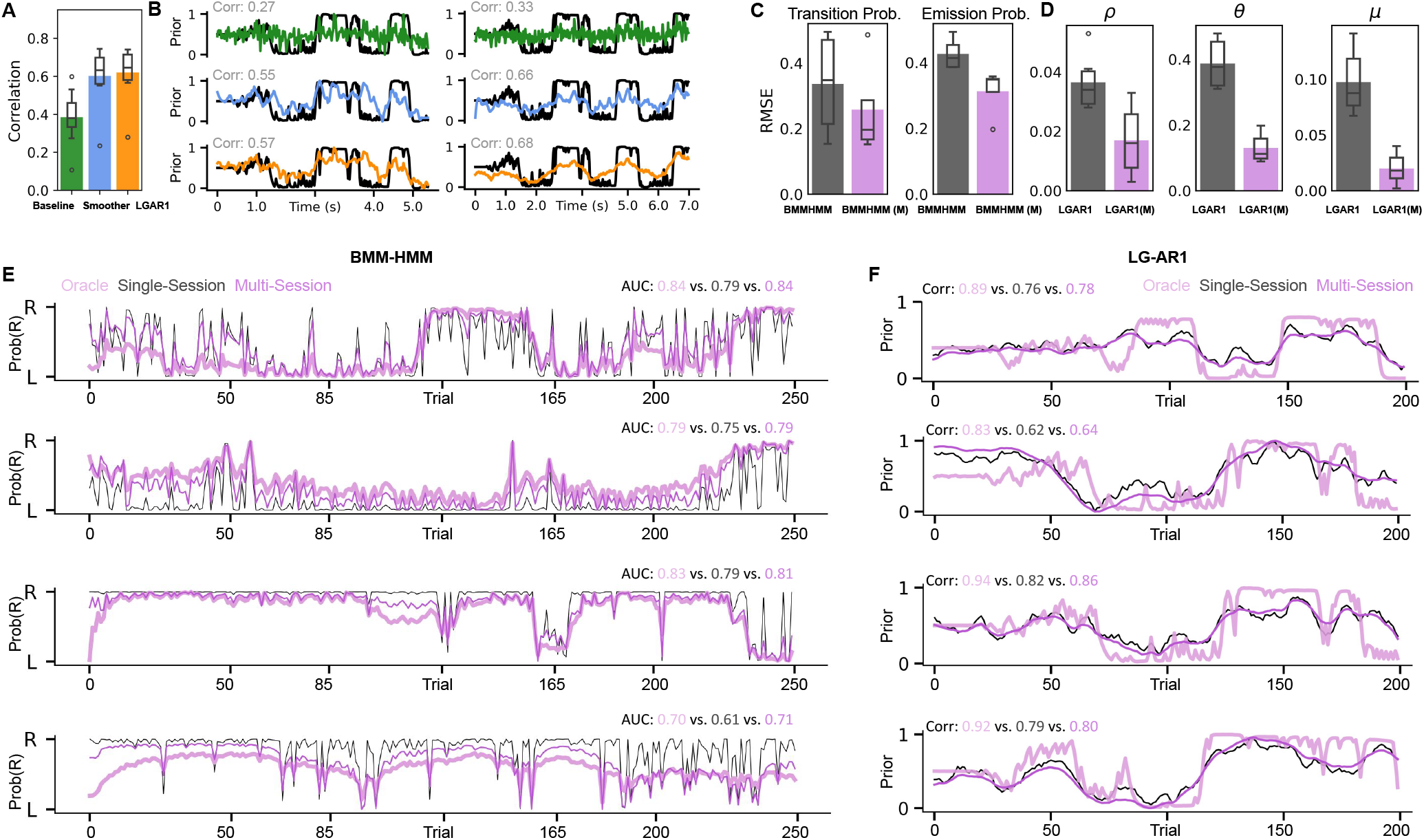
Relationship between LG-AR1 and post-hoc smoother, and across-session variability in parameter estimation and predictions of BMM-HMM/LG-AR1 models. **(A)** Comparison of prior decoding performance, measured by correlation, across the linear baseline, single-session LG-AR1, and a post-hoc smoother. Box plots display the minimum, maximum, median, and quartiles over 10 sessions, while bar plots show the mean correlation. **(B)** Predicted priors from the linear baseline, single-session LG-AR1, and post-hoc smoother are compared to ground truth in two example sessions. **(C-D)** Comparison of model parameter estimation accuracy, measured by root mean squared error (RMSE), between single-session models (BMMHMM, LGAR1) and multi-session models (BMMHMM (M), LGAR1 (M)). Box plots show the minimum, maximum, median, and quartiles across 10 sessions, while bar plots show the mean RMSE. For the BMM-HMM, RMSEs for the transition and emission probabilities are computed for each matrix entry in Figure 4E and then averaged. **(E)** Additional examples of decoded right-choice probabilities from the oracle (pink), single-session (black), and multi-session (purple) BMM-HMM models across 4 selected sessions. **(F)** Additional examples of decoded priors from the oracle (pink), single-session (black), and multi-session (purple) LG-AR1 models across 4 selected sessions.

**Figure S6:**
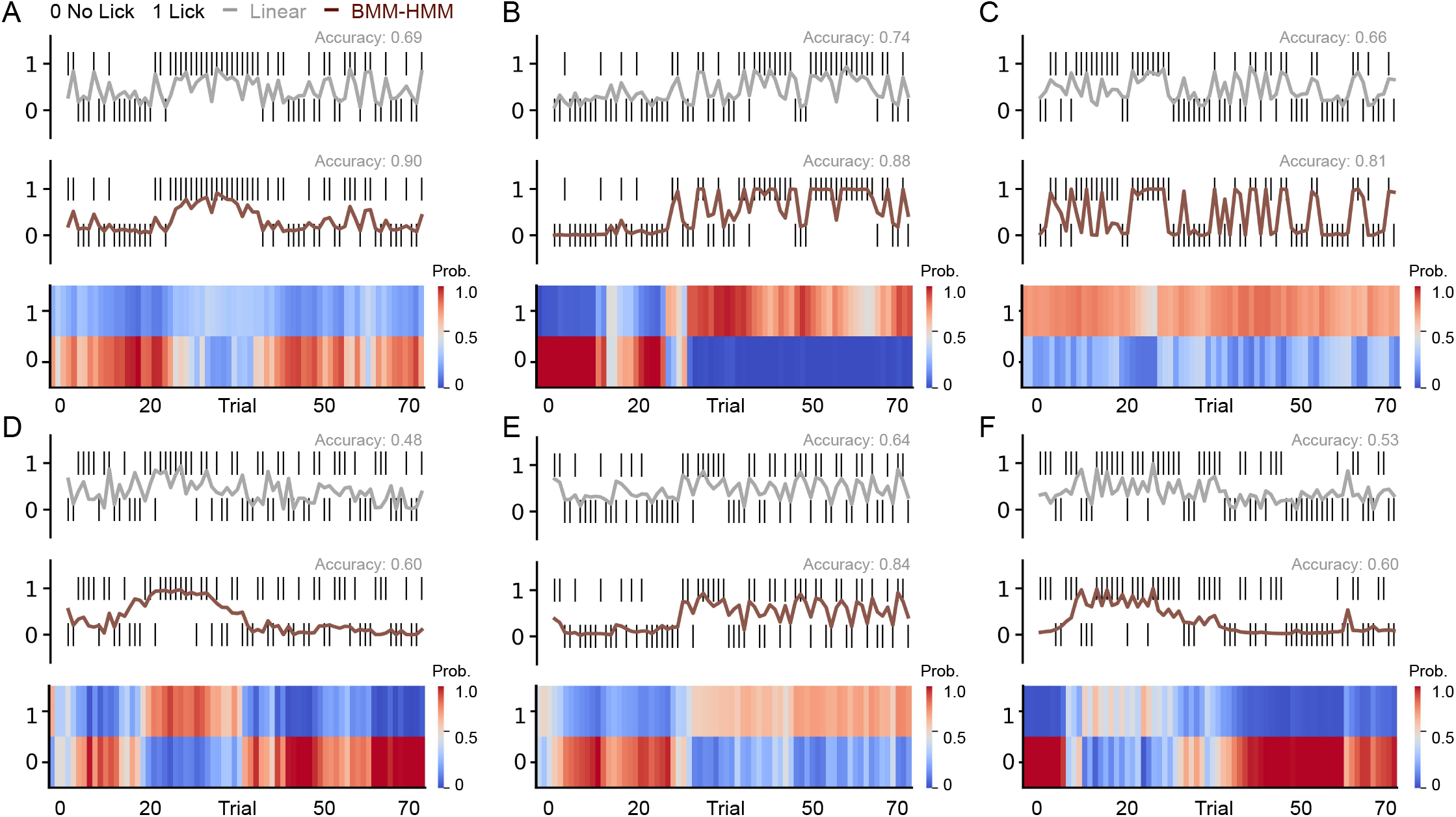
Additional examples of using BMM-HMM to infer latent states in licking behavior for better predictions in Allen datasets. **(A-F)** Comparison of the BMM-HMM model and a linear baseline in predicting behavioral responses (licks) to visual stimulus changes across six sessions, complementing the example shown in Figure 8I. For each session, we decode using neural activity from a subset of brain regions (DG, CA1, CA3, VISp, VISa, LP and TH), depending on which regions were recorded in that session. The inferred latent states are visualized as heatmaps, color-coded by the probability of each state.

**Figure S7:**
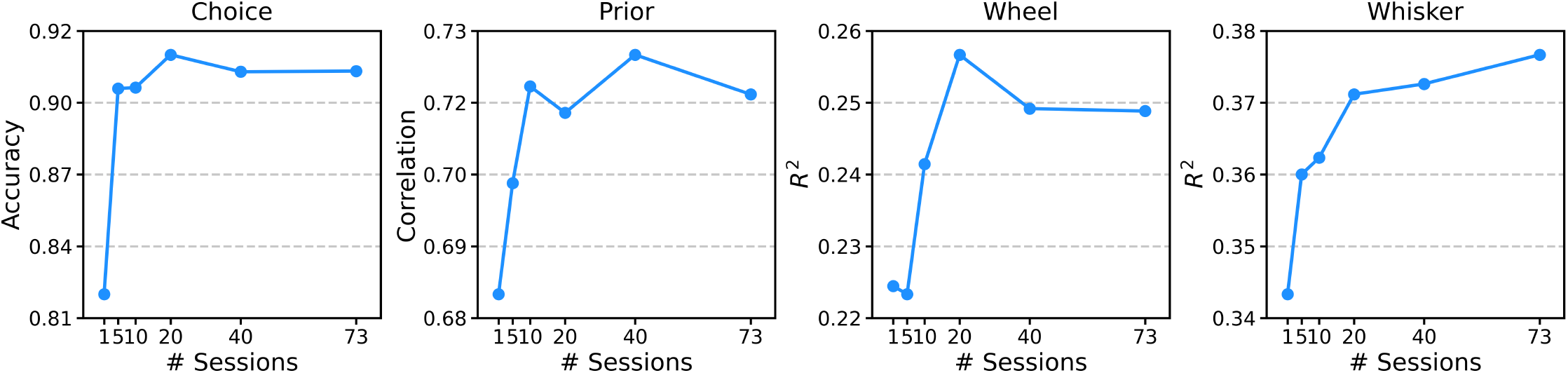
Scaling curves of multi-session reduced-rank regression models. For each behavioral variable, we train both single-session and multi-session reduced-rank regression models using data from 5, 10, 20, 40, and 73 sessions, and evaluate performance on a fixed set of 5 test sessions. Each panel shows how the average decoding score across these test sessions changes with the amount of training data.

