## Supplemental Tables and Figures for "Exploiting correlations across trials and behavioral sessions to improve neural decoding": Supplementary Materials.pdf

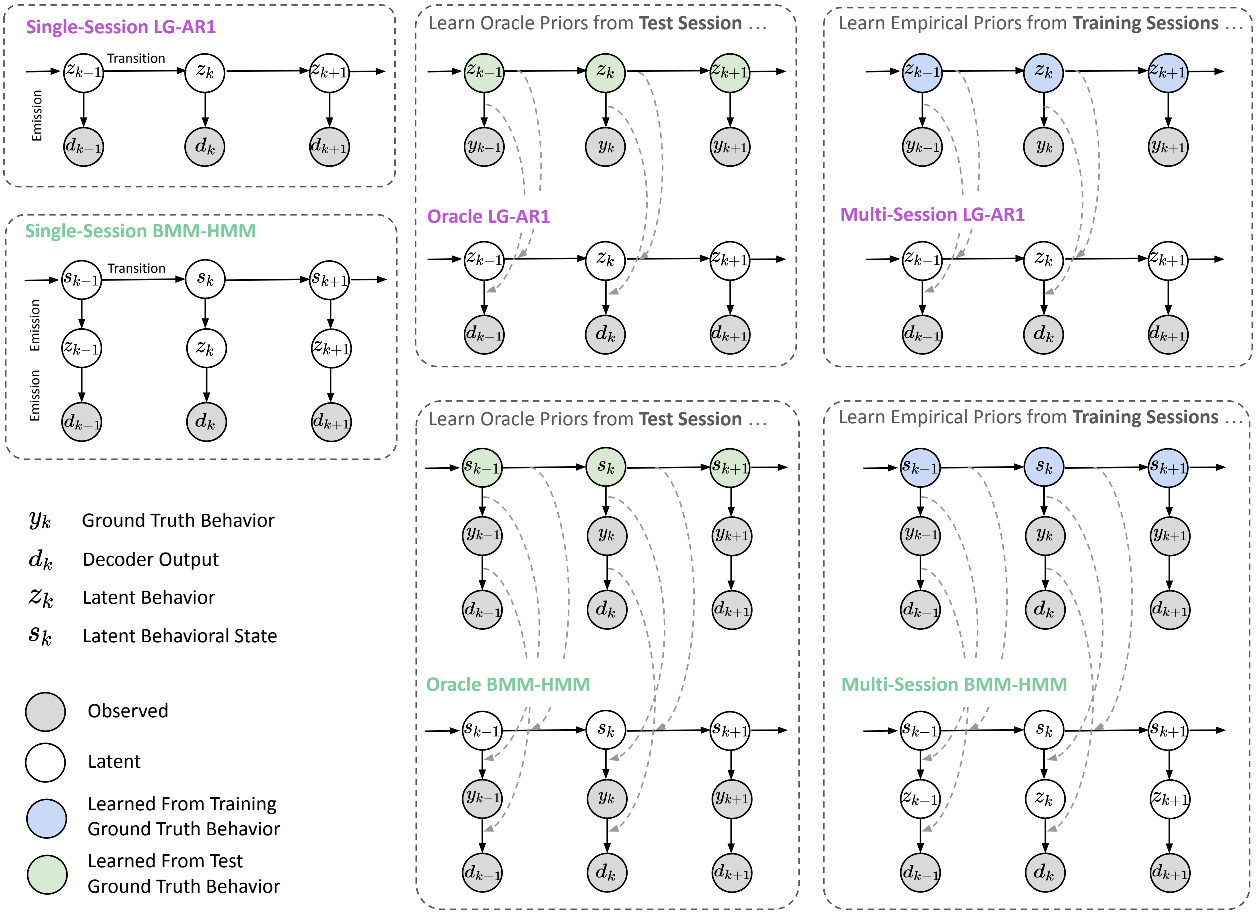


**Figure S1. Graphical models for single-session, oracle, and multi-session LG-AR1 and BMM-HMM models.** The single-session LG-AR1 and BMM-HMM models learn parameters directly from the test session. In contrast, the oracle LG-AR1 and BMM-HMM use a two-step learning process. They first derive oracle priors from the ground truth behavior in the test set, and then use these priors to guide parameter learning on the test data (shown by gray arrow). The multi-session LG-AR1 and BMM-HMM follow a similar approach to learn empirical priors from ground truth behaviors in the training data, which are then used to constrain parameter updates on the test set (shown by gray arrow). This approach allows the multi-session models to utilize information from multiple training sessions while adapting to the current test data.

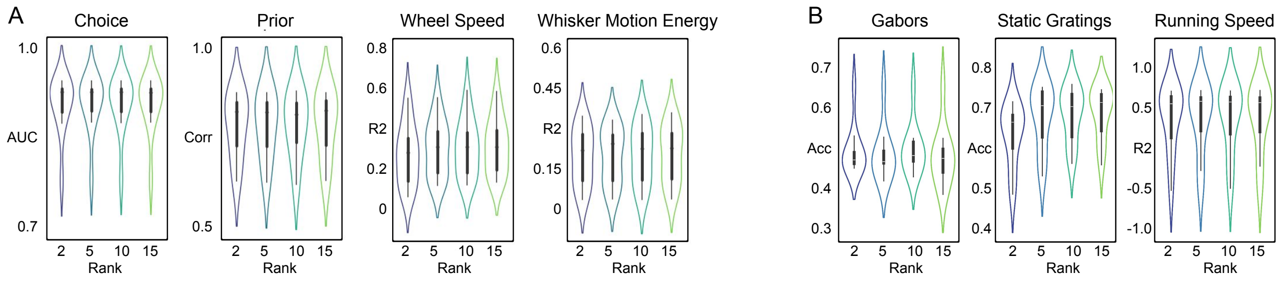


**Figure S2. Effects of model rank on decoding performance across different behavior variables.** For each behavior task in the IBL (A) and Allen (B) datasets, the violin plots display performance metric distributions across 10 sessions when decoding using different model ranks. Although all behaviors can be captured using lower model ranks, wheel speed and static gratings exhibit improved decoding performance at higher ranks.

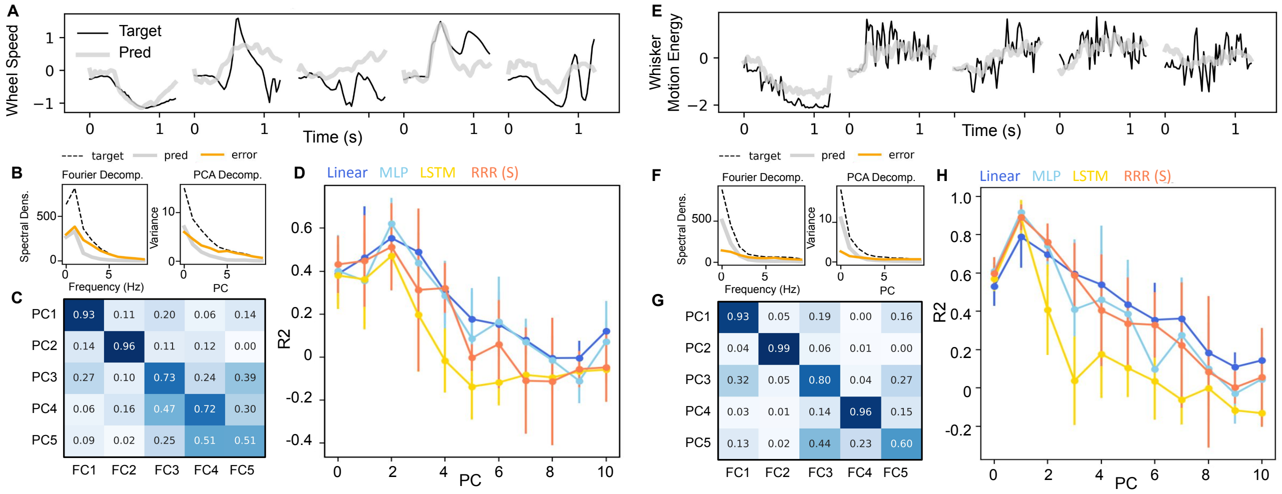


**Figure S3. Evaluating the reduced-rank regression model against baseline decoders in capturing the primary components of the target behaviors.**
Examples of actual (“Target”) and predicted (“Pred”) behaviors for wheel speed (A) and whisker motion energy (E) from the reduced-rank regression model across five selected trials. Whisker motion energy exhibits higher-frequency fluctuations compared to wheel speed.
Power spectral density (PSD) as a function of frequency is shown for the observed (“target”) and predicted (“pred”) wheel speed (B) and whisker motion energy (F) from the reduced-rank regression model, and decoding error (“error,” defined as target minus pred), averaged over 10 sessions. In addition, the variance explained by each principal component (PC) is plotted for the target, pred, and error. Both the low-frequency Fourier components and leading PCs account for most of the variance in the behavior signals.
Heatmap showing the Pearson correlation between the first 5 PCs and the first 5 Fourier components (FCs) of wheel speed (C) and whisker motion energy (G). A clear one-to-one correspondence is observed for the first two PCs and FCs, indicating that the leading PCs capture low-frequency components in the behavior.
Model performance ($R^2$) for decoding wheel speed (D) and whisker motion energy (H) reconstructed from each PC of the observed behavior using baseline decoders. Decoding $R^2$ is generally higher for the leading PCs, which are associated with low-frequency Fourier components. The mean and standard deviation of $R^2$ across 10 sessions are shown for the first 10 PCs.

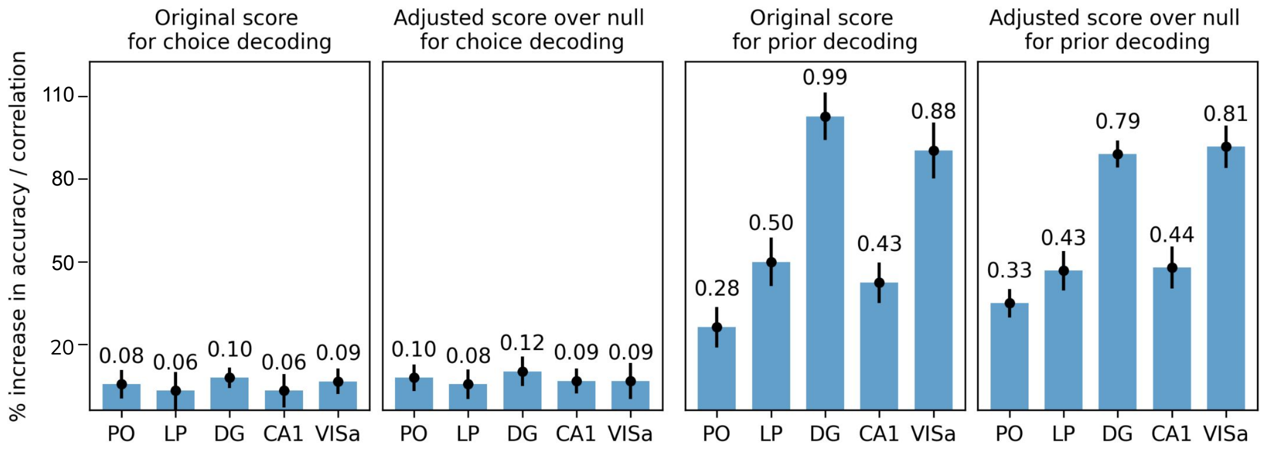


**Figure S4. Assessing the decoding improvement achieved by multi-region reduced-rank regression model relative to null distributions generated from imposter sessions.**
For each session with probe insertions in PO, LP, DG, CA1, and VISa, we create 10 “imposter sessions” from behaviors (choice and prior) of other mice in different IBL sessions. These are generated by concatenating trials across all analyzed sessions, excluding the session under consideration, then randomly selecting a chunk of $N$ consecutive trials (where $N$ matches the original session length) from the concatenated sessions. We obtain the original score from the real session, while the adjusted score is calculated by subtracting the decoding accuracy (or correlation) of the imposter sessions from the original score. Each bar shows the mean score from 10 imposter sessions, with error bars indicating one standard deviation of these scores.

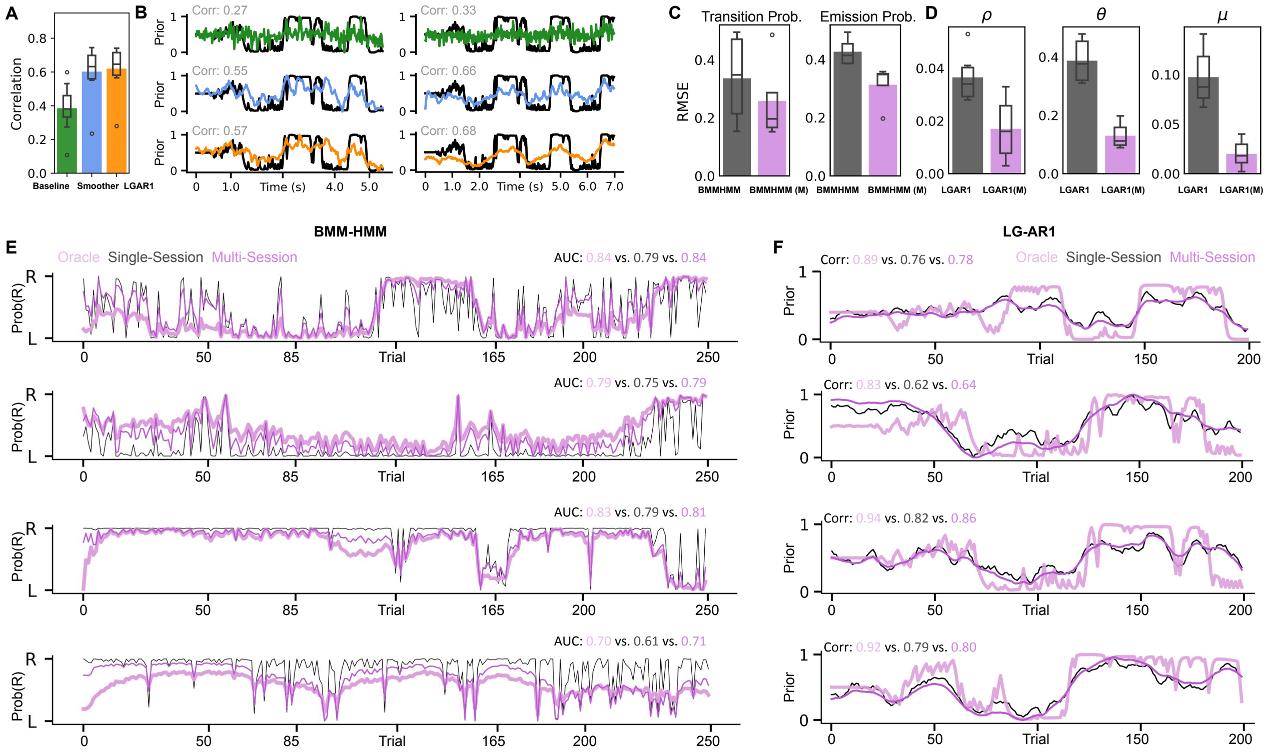


**Figure S5. Relationship between LG-AR1 and post-hoc smoother, and across-session variability in parameter estimation and predictions of BMM-HMM/LG-AR1 models.**

(A) Comparison of prior decoding performance, measured by correlation, across the linear baseline, single-session LG-AR1, and a post-hoc smoother. Box plots display the minimum, maximum, median, and quartiles over 10 sessions, while bar plots show the mean correlation.

(B) Predicted priors from the linear baseline, single-session LG-AR1, and post-hoc smoother are compared to ground truth in two example sessions.

Comparison of parameter estimation accuracy for BMM-HMM (C) and LG-AR1 (D) models, measured by root mean squared error (RMSE), between single-session models (BMMHMM, LGAR1) and multi-session models (BMMHMM (M), LGAR1 (M)). Box plots show the minimum, maximum, median, and quartiles across 10 sessions, while bar plots show the mean RMSE. For the BMM-HMM, RMSEs for the transition and emission probabilities are computed for each matrix entry in Figure 4E and then averaged.

(E) Additional examples of decoded right-choice probabilities from the oracle (pink), single-session (black), and multi-session (purple) BMM-HMM models across 4 selected sessions.

(F) Additional examples of decoded priors from the oracle (pink), single-session (black), and multi-session (purple) LG-AR1 models across 4 selected sessions.

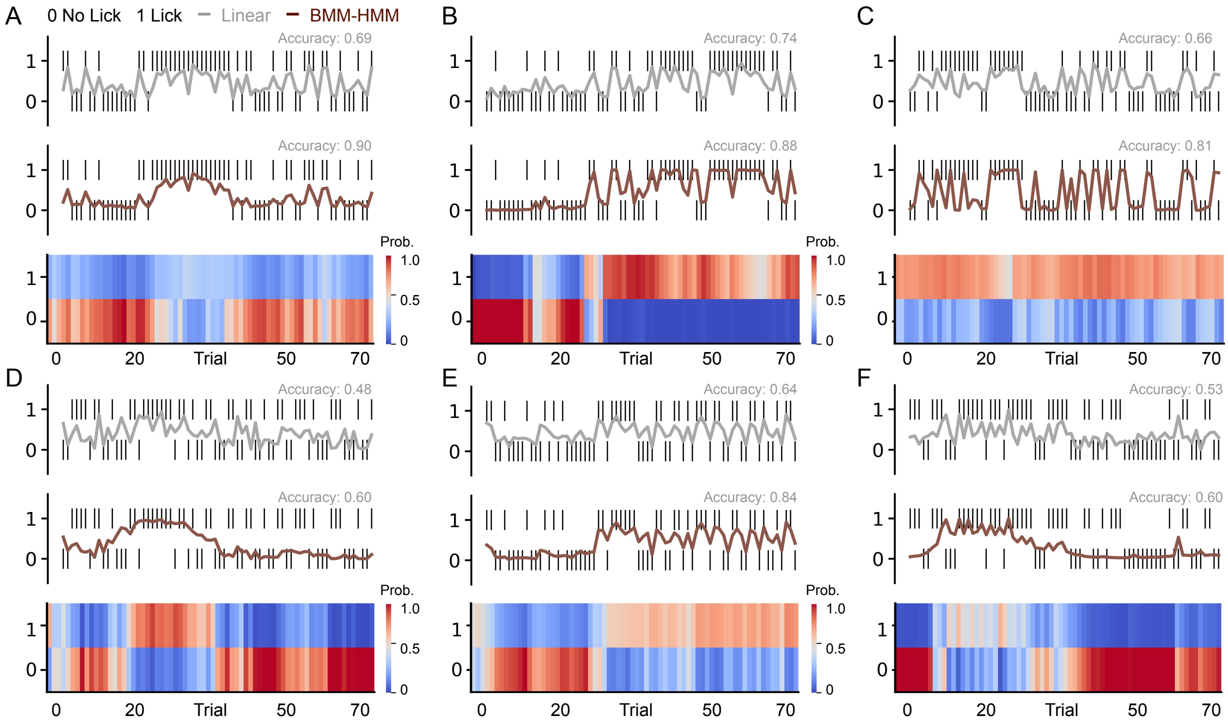


**Figure S6. Additional examples of using BMM-HMM to infer latent states in licking behavior for better predictions in Allen datasets.**
Comparison of the BMM-HMM model and a linear baseline in predicting behavioral responses (licks) to visual stimulus changes across six sessions, complementing the example shown in Figure 8I. For each session, we decode using neural activity from a subset of brain regions (DG, CA1, CA3, VISp, VISa, LP and TH), depending on which regions were recorded in that session. The inferred latent states are visualized as heatmaps, color-coded by the probability of each state.

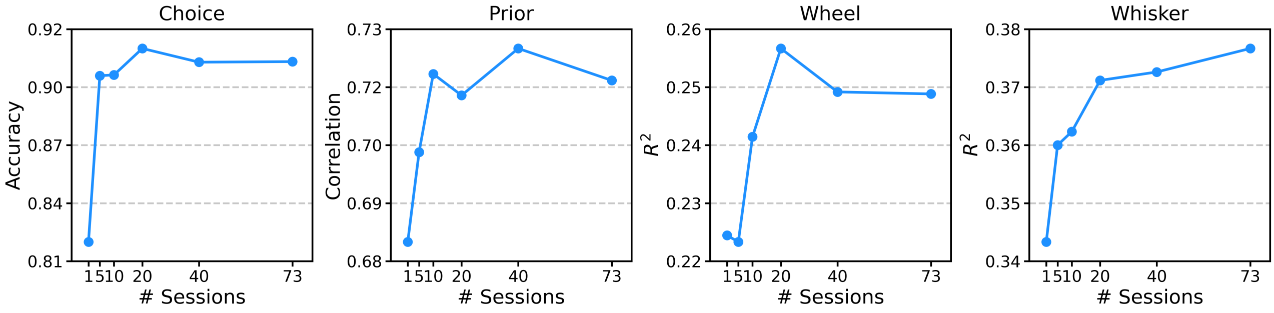


**Figure S7. Scaling curves of multi-session reduced-rank regression models.**
For each behavioral variable, we train both single-session and multi-session reduced-rank regression models using data from 5, 10, 20, 40, and 73 sessions, and evaluate performance on a fixed set of 5 test sessions. Each panel shows how the average decoding score across these test sessions changes with the amount of training data.

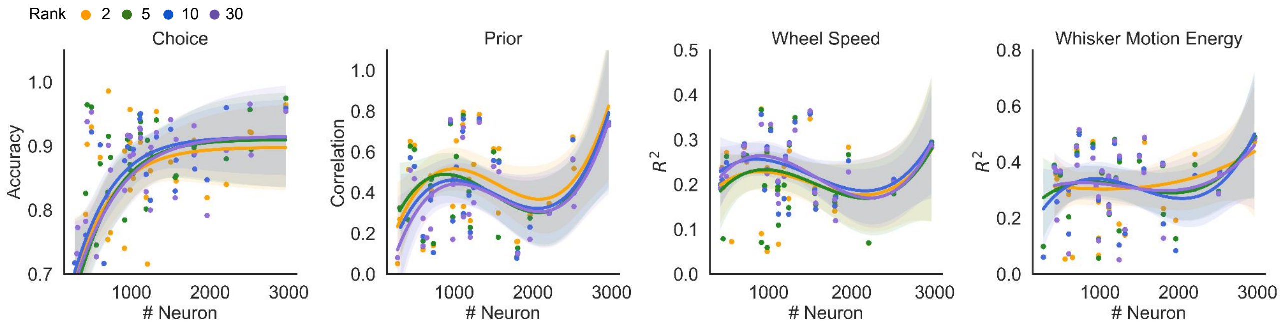


**Figure S8. Performance of single-session reduced-rank regression (RRR) model as a function of neuron count and model rank.**
We train single-session RRR on 48 IBL sessions with varying neuron counts to predict choice, prior, wheel speed, and whisker motion energy. Decoding performance is plotted against neuron count after excluding sessions with outlier metrics, with each dot representing one session. Colors distinguish different model ranks. To visualize the trends, we fit a logistic regression for choice and a third-order polynomial regression for the other tasks. Shaded areas denote the 95% confidence interval. For choice decoding, rank 2 achieves the highest performance when the number of neurons is fewer than 1500. For prior decoding, rank 2 is consistently the best across all neuron counts. For whisker motion energy, rank 2 outperforms higher ranks when the neuron count exceeds 1500. For wheel speed, larger model ranks lead to better performance with lower neuron counts.

**Table S1. Mathematical notation.**

Throughout the main text, STAR Methods section, and supplemental information, we use the mathematical notation defined in Table 1.

| Model | Notation | Dimensionality | Definition |
|---|---|---|---|
| RRR | $X$ | *N x T* | Single-trial neural activity |
| | $y$ | *P* | Single-trial ground truth behavior |
| | $d$ | *P* | Single-trial predicted behavior (decoder estimate) |
| | $U$ | *N x R* | RRR model's neural basis set |
| | $V$ | *R x T x P* | RRR model's temporal basis set |
| | $b$ | *P* | RRR model's intercept term |
| | $A$ | *L x R* | Multi-region RRR model's neural basis set for each brain region |
| | $B$ | *L x T x P* | Multi-region RRR model's temporal basis set shared across all regions |
| | $N$ | – | Number of neurons in a session |
| | $T$ | – | Number of time bins in each trial |
| | $K$ | – | Number of trials in a session |
| | $P$ | – | Dimension of the behavior of interest |
| | $R$ | – | Rank of the (multi-session) RRR model's $U$ and $V$ bases |
| | $L$ | – | Rank of the multi-region RRR model's $A$ and $B$ bases |
| BMM-HMM | $y_k$ | 1 | True behavior in trial *k* |
| | $d_k$ | 1 | Single-trial, single-session decoder estimate in trial *k* |
| | $z_k$ | 2 | Latent mixture assignment of trial *k* in a beta-mixture model |
| | $s_k$ | *H* | Hidden Markov model's latent state in trial *k* |
| | $\alpha_k(h)$ | 1 | Probability of past observations $d_1, \ldots, d_k$ at state *h* in trial *k* |
| | $\beta_k(h)$ | 1 | Probability of future observations $d_{k+1}, \ldots, d_K$ at state h in trial *k* |
| | $\gamma_k(h, y)$ | 1 | Probability of *y* at state *h* in trial *k* given $d_1, \ldots, d_K$ |
| | $\xi(h, m)$ | 1 | Transition probability from state *h* in trial *k* to state *m* in trial *k + 1* |
| | $\pi$ | *H* | HMM's initial state distribution |
| | $\eta$ | *H x H* | HMM's transition probability matrix |
| | $\phi$ | *H x 2* | HMM's emission probability matrix |

| | | | |
|---|---|---|---|
| | *H* | – | Number of latent states in an HMM |
| LG-AR1 | $y_k$ | 1 | True behavior in trial *k* |
| | $d_k$ | 1 | Single-trial, single-session decoder estimate in trial *k* |
| | $z_k$ | 1 | LG-AR1's latent state in trial *k* |
| | $\tilde{d}_k$ | 1 | Improved decoder estimate in trial *k* given $d_1, \ldots, d_K$ |
| | $\theta$ | 1 | LG-AR1's observation model parameter |
| | $\rho$ | 1 | LG-AR1's dynamic model parameter |
| | $\mu$ | 1 | Intercept term of LG-AR1's observation model |
| | $\sigma_\epsilon^2, \sigma_\tau^2$ | 1 | Variance of LG-AR1's noise term |
| | $\Lambda$ | – | LG-AR1 model parameters ($\theta$, $\rho$, $\mu$, $\sigma_\epsilon^2, \sigma_\tau^2$) |

**Table S2. The range of possible model and optimizer hyperparameters from which *Ray Tune* randomly samples combinations.**

| **Hyperparameter** | **Value Range** |
|---|---|
| RRR Rank | Randint(2, 50) |
| MLP Depth | Randint(1, 6) |
| MLP Hidden Size | [16, 32, 64, 128, 256, 512] |
| MLP Dropout Ratio | Uniform(0.1, 0.3) |
| Weight Decay | Log-Uniform(0.001, 1) |
| Learning Rate | Log-Uniform(0.0001, 0.01) |

**Table S3. Computation time for single- and multi-session reduced-rank regression (RRR) models and MLPs.** For single-session models, we report the mean and standard error of training times (in seconds) across 58 individual sessions. For multi-session models, we report the total time required to train a single model on all 58 sessions.

| **Time (s)** | **RRR** | **MLP** |
|---|---|---|
| Single Session | 129.10 $\pm$ 4.33 | 297.91$\pm$ 29.14 |
| 58 Session | 6610.70 | 16526.76 |
